# Synaptic input architecture of dendrites of V1 neurons revealed by large-scale synapse imaging without backpropagating action potentials

**DOI:** 10.1101/2024.10.24.619996

**Authors:** Satoru Kondo, Kohei Kikuta, Kenichi Ohki

## Abstract

How neurons integrate thousands of synaptic inputs to compute sharply tuned outputs is a critical question in sensory information processing. To answer this question, it is essential to record the location and activity of synaptic inputs *in vivo*. However, back-propagating action potential (BAP) calcium signals invade dendrites and spines, making accurate recording of spine responses difficult. In this study, we first developed a new method to record spine calcium responses without BAP signals. Using this method, we performed large-scale imaging of visually evoked spine activity from layer 2/3 pyramidal neurons and revealed three patterns of dendritic functional architectures of synaptic inputs: dendrites with clusters of spines of similar responses, dendrites with spines of diverse responses, and dendrites with spines where the majority of them show no visual response. Our model suggests that only a small fraction of spines on dendrites of clustered architectures are sufficient to generate sharply tuned output.

## INTRODUCTION

A single neuron in the central nervous system receives thousands of synaptic inputs and integrates them to generate an action potential (**Stuart & Spruston, 2015**). Because the depolarization induced by each individual synapse is relatively weak (**Mitchell et al., 2019**), the mechanism by which a single neuron determines its specific output from multiple inputs is not well understood. Over the past few decades, the morphology and electrical properties of dendrites have been studied intensively (**Stuart & Spruston, 2019**). Clustered synaptic activity in dendrites could lead to enhanced dendritic responses, potentially important for dendritic computation, was first proposed by a modeling study (**Mel, 1992**). Later studies have extensively explored this proposal through computer simulations (**Hausser & Mel, 2003**) and experimental methods by electrophysiology or imaging (**Magee, 2000; Losonczy & Magee, 2006; Losonczy et al., 2008; Branco & Hausser, 2010**). Furthermore, the somatic action potentials occur when the membrane potential depolarized by the sum of synaptic activities is only above the threshold (**Golding & Spruston, 1998**). Thus, these studies suggest the importance of both simultaneous clustered synaptic activation and the generation of above-threshold somatic action potentials for output signal generation. Because electrophysiological recordings of dendrites do not have sufficient spatial resolution for individual synaptic inputs, it is essential to record both the location and activity of individual synaptic inputs along dendrites using other techniques to answer the question of how neurons integrate synaptic inputs and determine output signals.

Recent technical advances in optical imaging, in particular two-photon calcium imaging, have made it possible to record brain activity in living animals with high spatial and temporal resolution (**Helmchen & Denk, 2005**). Using this technique, a large number of synaptic input signals distributed over dendrites can be measured with a high spatial resolution (**Jia et al., 2010; Chen et al., 2013; Grienberger et al., 2015**). In the case of pyramidal neurons, excitatory synaptic inputs mainly occur at small dendritic protrusions called spines (**Harris & Kater, 1994; Knott et al., 2006; Iascone et al., 2020**). Activation of a single spine induces an increase in calcium transients mediated by N-methyl-D-aspartate (NMDA) receptors, voltage-gated calcium channels or internal calcium stores (**Grienberger & Konnerth, 2012; Rochefort & Konnerth, 2012**) and this activity can be measured by calcium indicators (**Jia et al., 2010; Chen et al., 2011; Chen et al., 2013**). Using two-photon calcium imaging, the input-output transformation principle of a single neuron has recently been investigated, particularly with regard to neuronal response selectivity in V1 neurons (**Chen et al., 2013; Wilson et al., 2016; Iacaruso et al., 2017; Scholl et al., 2021**). The computation of neuronal response selectivity in V1 may provide a general model for understanding the input-output transformation principle of cortical neurons.

A prominent feature of V1 excitatory neurons is orientation selectivity, characterized by selective responses to a particular orientation of lines (**Hubel & Wiesel, 1962**). In higher mammals such as monkeys, cats and ferrets with functional columns, these orientation-selective neurons are arranged in a columnar structure, called orientation column. Animals with such functional column, a single excitatory neuron receives relatively uniform inputs from other excitatory neurons belonging to the same orientation-preference column and the mechanism to generate sharp outputs from uniform inputs has been explained experimentally in ferrets (**Wilson et al., 2016**). By contrast, in mice without functional columns (salt-and-pepper structure; **Ohki et al., 2005; Ringach et al., 2016; Kondo et al., 2016**), a single excitatory neuron generally receives diverse inputs from surrounding excitatory neurons with diverse responses. The predicted output signal from the simple summation of total inputs has much broader tuning than the soma (Chen et al., 2013; Scholl et al., 2021), and the mechanism that generates the sharp tuning of the soma remains unknown.

Several mechanisms for the generation of sharply tuned outputs have been previously reported. In ferrets, Wilson et al. suggested a contribution of dendritic nonlinearity for the sharpening of the tuning elicited by clustered synaptic activation (**Wilson et al., 2016**). This sharpening mechanism is supported by a study showing that visually evoked NMDA spikes on dendrites enhanced soma orientation selectivity in mice (**Smith et al., 2013**). Another possible mechanism for sharpening is thresholding at the soma. Somatic spiking is regulated by membrane potential thresholding, which allows firing in response to the preferred orientation and prevents firing in response to a non-preferred orientation (**Priebe & Ferster, 2008**). Electrophysiological recordings from cat V1 neurons showed that the orientation tuning of spike responses was narrower than the tuning of the membrane potential (**Carandini & Ferster, 2000**). In contrast, simultaneous *in vivo* spine imaging and electrophysiological recording of soma membrane potential showed a highly variable relationship between spine activity and somatic firing (**Wilson et al., 2016**).

To elucidate the mechanisms to generate the sharp tuning in mice, we first developed a novel method that solves the critical problem in *in vivo* spine imaging: the intrusion of back propagating action potential (BAP) calcium signals into the spine, which potentially leads to inaccurate measurements of the spine calcium signal. The mathematical method has been used to isolate the spine calcium signal by subtracting the dendritic calcium signal (**Chen et al., 2013; Wilson et al., 2016; Iacaruso et al., 2017; Scholl et al., 2021**) but the question remains how accurately this method estimates true spine activity.

In this study, we established a new optogenetic method for imaging spine activity without BAPs using a bistable inhibitory opsin containing a soma-targeting signal. This method has a major advantage over previously reported BAP-inhibition methods (**Jia et al., 2010; Levy et al., 2012**) in that the prevention of somatic depolarization by photoinhibition is technically easier and lasts for longer, and BAP-free imaging can be performed during this period. By expressing a bistable inhibitory opsin, we could suppress the soma activity effectively and successfully separate the spine signals from the BAP signals.

Using this method, we conducted large-scale spine imaging (∼1,000 spines per neuron) from layer 2/3 pyramidal neurons in mouse primary visual cortex and obtained new findings. First, functional architecture of dendrites can be classified into three synaptic input patterns.: dendrites with clusters of spines of similar responses, dendrites with spines of diverse responses, and dendrites with spines where the majority of them show no visual response. Next, to predict the sharp output tuning, we examined several theoretical models based on our experimental data and found that the dendritic cluster model, which only considers the dendrites with cluster of spines of similar responses for the control of output signal, is effective in predicting the sharp output tuning. Our data suggest that only a small fraction of spines (6.9% of the total spines) form clusters of similar inputs on a small subset of dendrites (21.9% of total dendrites), and this functional architecture effectively contributes to the generation of sharply tuned output.

## RESULTS

### Prevention of BAP by inhibitory opsin

First, to analyze how strongly the synaptic input signals are affected by BAP signal along dendrites, we sparsely expressed (**Xu et al., 2012**) the genetically encoded calcium indicator GCaMP6s in layer 2/3 pyramidal neurons (see Methods) and imaged the activity of individual spines in response to drifting gratings (Figure S1). The expression of GCaMP6s was sufficiently sparse, making it unlikely that the spines we imaged belonged to neurons other than those subjected to somatic inhibition (Figure S1d, two-dimensional image perpendicular to the cortical surface; Figure S1e, three-dimensional image reconstructed from a stack of two-dimensional images). GCaMP6s was expressed throughout the dendrites and spines without filling the nucleus of the soma (Figure 1b, Figure S3a). The time course of calcium signal changes in dendrites closely resembled those of soma (Figure S1c, see colored single traces), suggesting that action potentials propagated back to dendrites. We noticed that spine signals also resembled those of soma and dendrites, suggesting that BAP signals invade individual spines and mask the original spine signals (Figure S1c, colored single traces).

**Figure 1.**
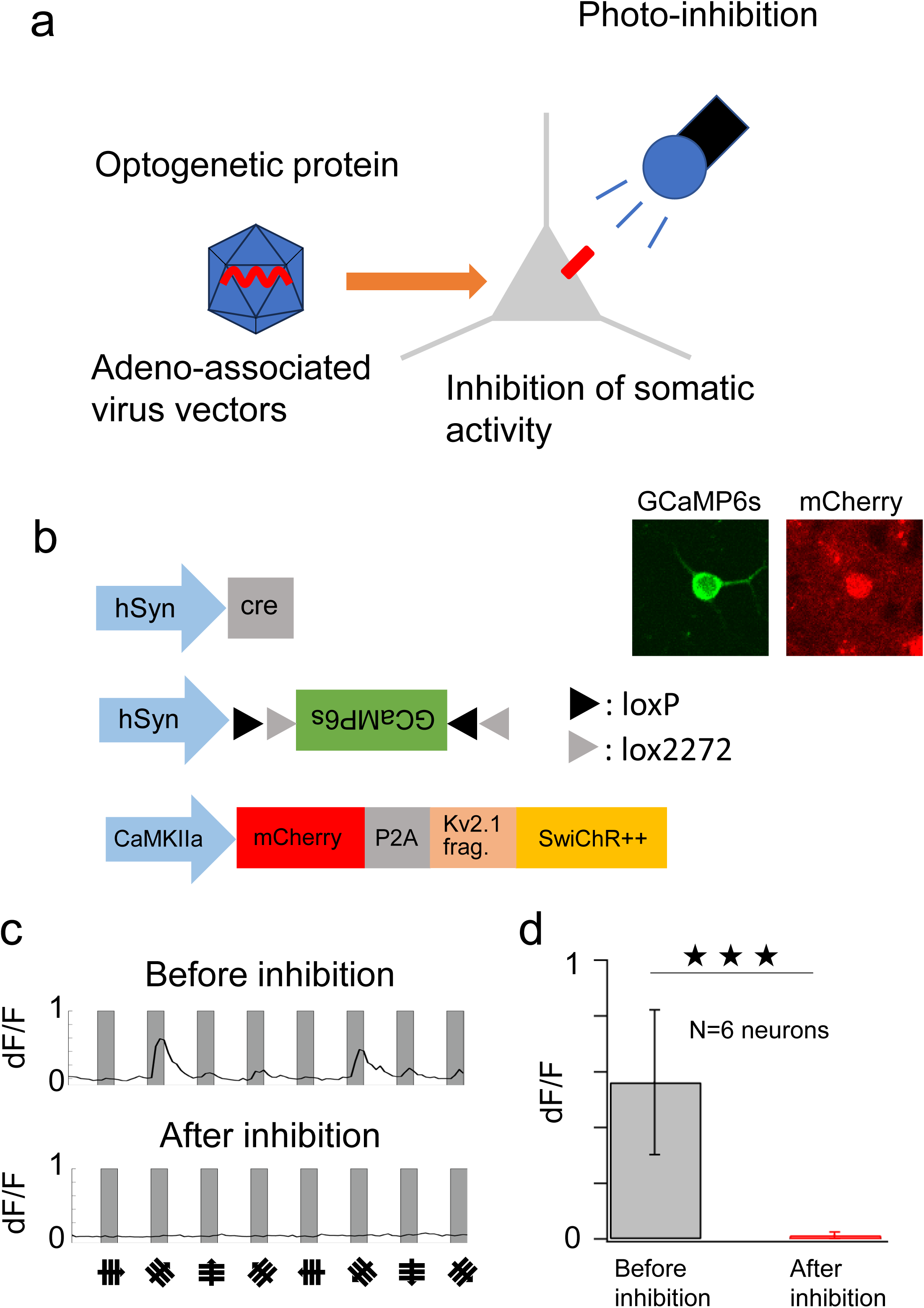
New method to record visual response of spines without BAP signals. **a.** A gene encoding the inhibitory optogenetic protein SwiChR++ is introduced by AAV into layer 2/3 pyramidal neurons in mouse V1. **b.** GCaMP6s is used for the functional spine imaging. Cre-loxP system is used for the sparse expression of GCaMP6s by diluting the Cre-expressing vector. SwiChR++ is used to inhibit the somatic activity. A mixture of three types of AAV is injected to V1 A neuron that expresses both GCaMP6s and mCherry which is the fluorescence marker for the expression of SwiChR++ is shown. **c.** Upper panel: Before photoinhibition, soma shows orientation-selective responses (p value for the response = 2.01E-8 and max ratio change=0.6057, see Methods for the definition). Lower panel: After photoinhibition, the visual response of soma is significantly abolished (p-value for the response = 0.097 and max ratio change=0.0129, see Methods for the definition). d. Averaged max ratio change from 6 neurons shows that the visual response of soma is significantly abolished (***: p<0.001, paired-sample t-test).

To prevent contamination of the BAP signal in the spine signal, we tried to suppress somatic activity using an inhibitory optogenetic method. To this end, we expressed the bi-stable (step-function) inhibitory opsin SwiChR++ (**Berndt et al., 2016**) in excitatory neurons. To confine the inhibitory effect to the soma, we added a soma-targeting signal to the opsin to localize its expression (**Lim et al., 2000**; Figure S2). We expressed GCaMP6s and SwiChR++ with adeno-associated virus (AAV) vectors using the Cre-loxP recombination system and photo inhibited somatic activity of neurons that co-expresses GCaMP6s and mCherry with a laser (Figure 1a, b). First, we attempted two-photon activation of SwiChR++ with pulsed lasers at wavelengths of 920 nm or 1,040 nm, but neither raster scans nor tornado scans restricted on the soma failed to suppress action potentials (data not shown). We next attempted one-photon activation of SwiChR++ using a visible light laser. Photoinhibition by a raster scan with a visible laser confined to the soma (wavelength: 458 nm, beam diameter: 0.53 μm, scan size: 20 x 20 pixels (0.5 μm/pixel), scan speed: 100 ms/frame, duration: 100 s, laser power: 50 μW after the objective lens) successfully prevented visually evoked somatic activity (Figure 1c, Before-inhibition: p value for the response = 2.01E-8, maximum response change = 0.6057, After-inhibition: p-value for the response = 0.097 and maximum ratio change = 0.0129 respectively (see Methods for the responsiveness criteria). To evaluate the effectiveness of the inhibition, we conducted a statistical analysis of the maximum ratio change in response to visual stimuli before (mean maximum response change = 0.562) and after BAP inhibition (mean maximum response change = 0.013) in six cells (n = 6 cells). A paired-sample t-test revealed a significant reduction in visual responses following BAP inhibition (p < 0.001), indicating the effectiveness of soma-targeted BAP inhibition (Figure 1d). The effect of one-time photoinhibition lasted for approximately an hour, and visually evoked spine activity could be measured without BAP contamination during this time period (Figure S3). By repeating the inhibition of the somatic activity, imaging can be extended to longer time period than previously reported BAP-inhibition methods, patch-clamping the cell body (**Jia et al., 2010; Chen et al., 2011**) or microinjecting a sodium channel blocker into the soma (**Levy et al., 2012**). Note that, the somatic tuning was measured before photoinhibition of the soma activity, so the measurement of the somatic tuning was not affected by photoinhibition.

### The validation of new technique to record spine activity without BAP signals

In order to validate our new technique for recording spine activity, we examined the effect of photoinhibition on the surrounding neurons of the targeted neuron and on the photoinhibited neurons’ own spine activity.

First, we investigated the impact of silencing a targeted neuron on the activity of neighboring neurons. Even if the light irradiation is restricted to a single neuron, this alteration has the potential to impact the circuit in which the neuron is embedded, especially its recurrent networks. We expressed a long-lasting inhibitory optogenetic protein (SwiChR++) in transgenic mice (Thy1-GCaMP6s, GP4.3) expressing GCaMP6s in the V1 and compared the response changes around the targeted neuron (250 μm radius) before and after photoinhibition. For the evaluation, we measured signal changes, preferred orientation and sharpness of orientation tuning (gOSI) as a function of distance from the targeted neuron at 3 different planes (light targeted plane and 30 μm upper and 30 μm lower to the light targeted plane). We observed statistically not significant response changes more than 50 μm away of the targeted neuron, but the response signal was slightly reduced in neurons within 50 μm radius of the targeted neuron (about 18% decrease on average; 368 orientation-selective neurons from 3 mice) on the same plane as the targeted plane (Figure S4, S5). Despite of the effect on signal changes, there was no significant difference in ΔOri (the difference of the preferred orientation between before and after inhibition) or gOSI even within 50 μm radius of the targeted neuron (Figure S6-S9). The same results were obtained on the different planes, upper and lower planes of the targeted planes (Figures S4, S6, S8; upper plane (+30 μm), 360 orientation-selective neurons from 3 mice; lower plane (−30 μm), 384 orientation-selective neurons from 3 mice).

Considering the effect of this small reduction (∼18%) of the response signal within 50 μm on synaptic inputs, previous studies examining the probability of connections between pyramidal neurons in layer 2/3 of the rat visual cortex showed that while the probability of connections between neurons within 50 μm was high, the absolute number of neurons connected was only about 16% of the total number of connected neurons (10 out of 61) due to the small spatial volume, and about 84% of neuronal connections (51 out of 61) were made with pyramidal neurons more than 50 μm apart (**Holmgren et al., 2003**). Therefore, it is likely that the slight suppression within 50 μm (∼18%) in the present study represents only a small proportion of the total input (∼16%) and the slight suppression of these neurons (∼18%) is unlikely to have a significant effect on synaptic inputs to the photoinhibited neuron as well.

Next, we examined the effect of soma silencing on the photoinhibited neurons’ own spine activities (Figure 2, Figure S11; 218 orientation-selective spines from 5 mice). Photoinhibition significantly suppressed action potentials and BAPs (Figure 1c). Spines whose preferred orientation is different from the soma are mostly unaffected by BAP. By comparing the response signals of these spines before photoinhibition (without subtraction) and after photoinhibition (BAP-inhibited), the effect of photoinhibition on the photoinhibited cell’s own spine can be evaluated. When comparing the spine response signals before and after photoinhibition, photoinhibition did not significantly affect the spine response signal (Figure 2c, p=0.86 paired sample t-test; Figure S11a, p=0.09 one sample t-test). These results indicate that somatic hyperpolarization does not significantly affect the photoinhibited cell’s own spine activities.

**Figure 2.**
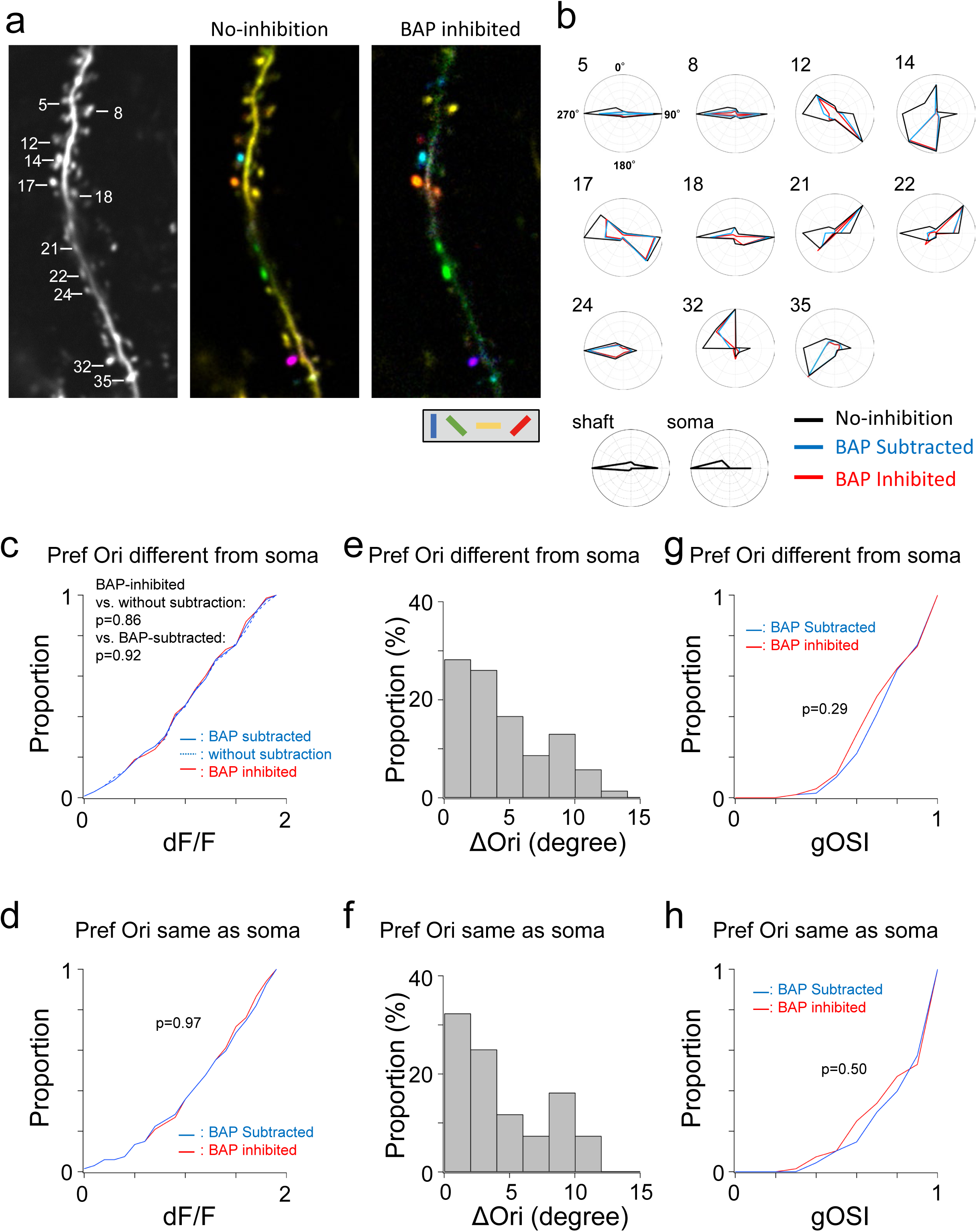
Comparison between the subtraction and photoinhibition methods for the visually responsive spine activity. The preferred orientation of the soma of this neuron is horizontal (coded yellow). **a**. Left: An example image of averaged dendritic branch. Middle: An orientation map of the same dendritic branch without BAP inhibition (no-inhibition). Right: An orientation map of the same dendritic branch after BAP inhibition (BAP inhibited). **b**. Polar plots of visual responses from orientation-selective spines. All the spines show similar orientation preference between BAP-subtracted and BAP-inhibited methods. Black: no-inhibition, Blue: BAP subtracted, Red: BAP inhibited. **c.** Cumulative plots of dF/F of all the orientation-selective spines with preferred orientation different from the soma of without subtraction, BAP-subtracted and BAP-inhibited methods. There is no statistically significant difference among different methods (p=0.86, without subtraction vs. BAP-inhibited; p=0.94, BAP-subtracted vs. BAP-inhibited; p=0.92, without subtraction vs. BAP-subtracted; paired-sample t-test). **d**. Cumulative plots of dF/F of all the orientation-selective spines with preferred orientation same as the soma of BAP-subtracted and BAP-inhibited methods. There is no statistically significant difference between two methods (p=0.45 paired-sample t-test). **e, f** Bar plots of difference of preferred orientation (ΔOri) between BAP-subtracted and BAP-inhibited spines with preferred orientation different from the soma (**e**) and same as the soma (**f**). **g.** Cumulative plots of gOSI of all the orientation-selective spines with preferred orientation different from the soma of BAP-subtracted and BAP-inhibited methods. There is no statistically significant difference between two-methods (p=0.29 paired-sample t-test). **h**. Cumulative plots of gOSI of all the orientation-selective spines with preferred orientation same as the soma of BAP-subtracted and BAP-inhibited methods. There is no statistically significant difference between two-methods (p=0.50 paired-sample t-test).

Then, we compared the two different methods, BAP-inhibition and BAP-subtraction methods. When comparing BAP-inhibited spine signals with BAP-subtracted spine signals, there was no statistically significant difference between BAP-inhibition and BAP-subtraction methods in spine response signals (Figure 2b-d, Figure S11a, b), preferred orientation (Figure 2e, f), or tuning sharpness (Figure 2g, h, Figure S11c, d). The same results were obtained for both types of spines whose preferred orientation is different from (not affected by BAP) (Figure 2c, p=0.92; e; g, p=0.29 paired sample t-test; Figure S11a, p=0.08; b, p<0.001 one sample t-test) or the same as (affected by BAP) (Figure 2d, p=0.45; e; h, p=0.50 paired sample t-test; Figure S11c, p=0.08; d, p<0.001 one-sample t-test) the soma.

These results indicate that somatic hyperpolarization does not affect the spine signal measurement in the photoinhibited cell, and the similar results were obtained in the spine signal measurements between the BAP-inhibited and BAP-subtracted methods. Previously, it was unknown whether the contaminated signal from BAP is linearly added to the spine signal and whether the contaminated signal can be accurately subtracted by a robust linear regression method (**Chen et al., 2013**; BAP subtraction method). The present study using the new BAP inhibition method clearly demonstrated the similar results between the two different methods. Our newly developed BAP inhibition method not only realized the measurement of the BAP-free spine signal for the first time, but also demonstrated the validity of the conventional BAP subtraction method, the accuracy of which in estimating the true spine signal was unknown.

### Functional input map of layer 2/3 pyramidal neurons in mouse V1

To understand how stimulus selectivity is computed in a single neuron, we reconstructed synaptic input maps from layer 2/3 pyramidal neurons in mouse V1 (Figure 3). To image spine activity, we used our new method to prevent somatic activity and recorded visually evoked spine signals under BAP-free experimental conditions. Among the recorded neurons (6 neurons, see Table 1), two neurons were orientation selective, one neuron was direction selective, and three neurons were both orientation and direction selective. We imaged the activity of ∼1,000 spines per neuron, and 21.6% (average from 6 neurons; range: 16.2-26.9%; Table 1) of the spines were significantly responsive (see Methods for the definition) to the drifting grating visual stimulus. Among the visually responsive spines, 85.4% (mean of 6 neurons; range: 82.1-90.0%; Table 1) of the spines showed orientation-selective responses, and 57.0% (mean of 6 neurons; range: 48.2-74.3%; Table 1) of the spines gave direction-selective responses (see Online Methods for the definition). The distributions of preferred orientation and direction of spines are shown in Figure 4a. As previously reported (**Chen et al., 2013; Wilson et al., 2016; Iacaruso et al., 2017; Scholl et al., 2021**), the proportion of spines with a preferred orientation similar to the soma (ΔOri <= 30°) was higher than the number with other orientations (60.0%, average of 6 cells; range: 44.7–77.0%; Figure 4b, c).

**Figure 3.**
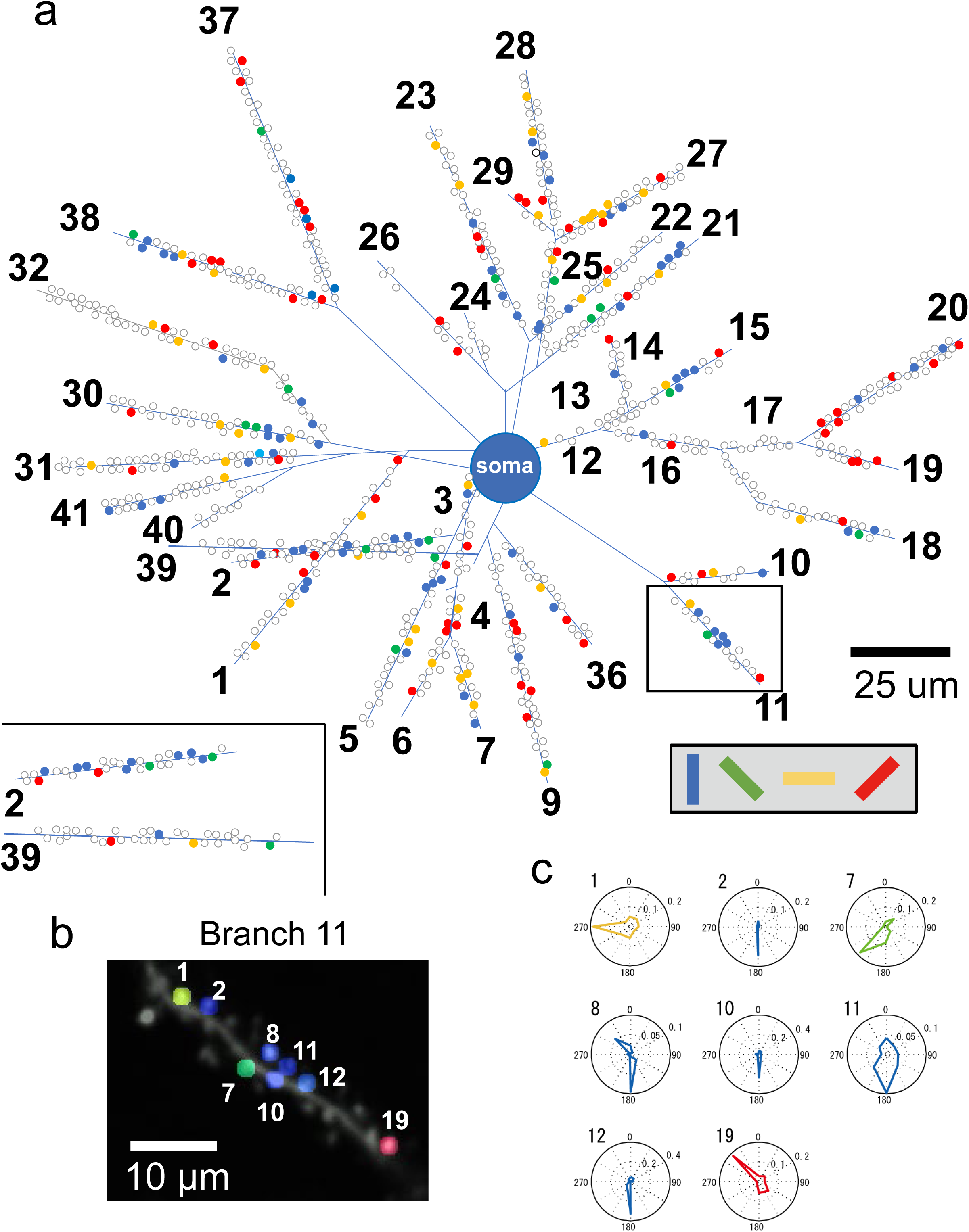
Functional spine map of a layer 2/3 pyramidal neuron. **a.** A functional spine map from ∼1,000 spines recorded from the basal dendrites of neuron 1. Orientation-selective spines are illustrated by different colors, and uncolored spines are either visually unresponsive or visually responsive but nonselective. **b.** A spine orientation map from branch 11 of neuron 1 (see Table 1). Among nineteen spines recorded from this branch, eight spines selectively responded to orientation stimulation. **c.** Polar plots of orientation-selective spines in branch 11.

**Table 1.** Summary of recorded spines.

| Neu<br>ron<br># | Soma<br>OSI | Soma<br>DSI | Recorded<br>spine N | N of<br>visually<br>responded<br>spine | N of<br>recorded<br>branch | N of<br>orientation-selective<br>spine | N of<br>direction-selective<br>spine |
| --- | --- | --- | --- | --- | --- | --- | --- |
| 1 | 1 | 0.45 | 996 | 261<br>(26.2%) | 41 | 234<br>(90.0%) | 194<br>(74.3%) |
| 2 | 1 | 0.51 | 920 | 217<br>(23.6%) | 32 | 188<br>(86.6%) | 119<br>(54.8%) |
| 3 | 1 | 0.01 | 723 | 117<br>(16.2%) | 24 | 96<br>(82.1%) | 62<br>(53.0%) |
| 4 | 1 | 0.25 | 993 | 198<br>(19.9%) | 41 | 165<br>(83.3%) | 115<br>(58.1%) |
| 5 | 0.32 | 0.65 | 1119 | 301<br>(26.9%) | 40 | 251<br>(83.4%) | 162<br>(53.8%) |
| 6 | 1 | 0.65 | 982 | 166<br>(16.9%) | 30 | 144<br>(86.7%) | 80<br>(48.2%) |

**Figure 4.**
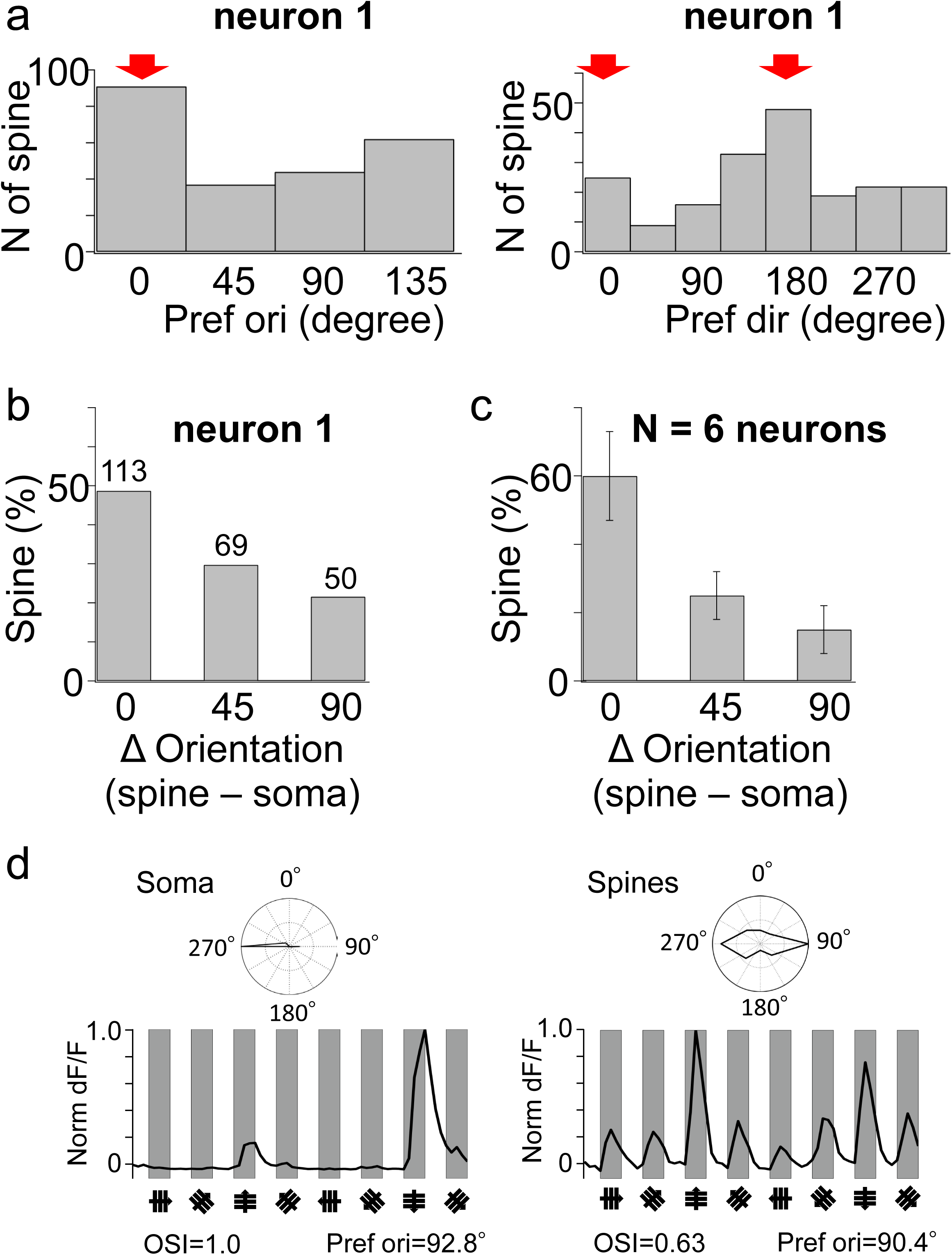
The preferred orientation of the soma can be predicted both from the number of orientation-selective spines and from the summed calcium time courses of spines, although with broader tuning. **a.** Distributions of preferred orientations (left) and directions (right) of spines in neuron 1. The peaks of the distribution match the preferred orientation and direction of the soma (red arrow). **b.** The distribution of the difference in preferred orientation between each spine and soma (ΔOrientation) of neuron 1. About half of spines are tuned similarly to the soma. **c.** Averaged data from 6 neurons shows the same trends as neuron 1 data. **d.** The time course and polar plot of the soma (left panel; average of 10 trials) and the average of the spine signals and polar plot (right panel; average of 10 trials from 982 spines). The OSI and preferred orientation are shown for both datasets. Polar plots suggest the broader tuning of the average of the spine signals than the tuning of the soma.

### Predicting somatic tuning from the number of responsive spines

To investigate whether the soma response could be estimated by linear summation of the spine responses, we first averaged the time courses (Figure 4d right) or tuning curves (data not shown) of the calcium signals from all recorded spines and compared them with the soma response. By averaging the signals from all spines, we estimated the orientation tuning curve of the spine inputs, which was able to predict the preferred orientation of the soma but was broader than that of the soma (Figure 4d). To assess the tuning sharpness and similarity between the signals of the soma and the averaged spines, we calculated the orientation selectivity index (OSI) and the correlation index (CI) (see Online Methods for the formula, CI = 1 means a perfect match between the two tuning curves). The OSI showed a much broader tuning in the averaged spines signal (OSI = 0.63) than in the soma signal (OSI = 1). The CI (CI = 0.69) indicated that the tuning curves between the soma and the averaged spines were not very similar. These results suggest that the simple summation of the spine responses is not a good predictor of somatic tuning.

The time course of signal changes in spines includes weak, subthreshold responses, and simple summations of spine responses may have broadened the tuning curve. To exclude the weak and subthreshold spine responses, instead of averaging the responses of all spines, we summed the number of all spines that significantly responded to the visual stimulation for each stimulus direction (Figures 6a, S11c). The polar plot showed that the tuning curve of the summed spines only slightly approached the tuning curve of the soma compared to the simple summation of the spine responses. When we compare the Figure 4d right (summation of dF/F) and Figure 6b (summation of number of responded spines), CI slightly increased from 0.69 (summation of dF/F) to 0.78 (summation of number of responded spines), and the preferred direction became the same as the soma (the preferred direction of Figure 6b is opposite to the soma), however the overall tuning curve of both are still dissimilar to that of soma. The increase of the response threshold did not further improve the CI (Figure S12a, d).

Since this summation is only a digital summation (0 or 1), we investigated incorporating a weighting for individual spines to improve the estimation of the somatic tuning. First, spine activity was weighted by the change in calcium signal (Figure 6c). The prediction of somatic tuning was not significantly improved compared to the result of unweighted summations (CI = 0.78 (unweighted) to 0.80 (weighted by calcium signal change)). Improvement of CI was not observed even if we increased the response threshold of summation (Figure S12b, d).

The effect of spine activity on the soma may depend on the distance of the spine from the soma, and the closer the spine is to the soma, the greater the effect (**Spruston, Jaffe & Johnston, 1994**). Therefore, we next considered the distance between the soma and the spines as a weighting factor. The weighting coefficient was defined as ‘exp (-distance/length constant)’, which is derived from the cable theory of dendrites (**Rall, 1959**). Based on the literature for layer 2/3 pyramidal neurons (**Larkum et al., 2007**), the length constant was set to 200 μm (Figure 6d). The predicted tuning curve was no better than without weighting (Figure 6b; CI = 0.78, both unweighted and weighted by distance from the soma). We also calculated the tuning curves while changing the length constant from 100 μm to 300 μm, but there was no difference in the tuning curves obtained (Figure S13a, b).

We also tested the possibility that spines located within a limited distance from the soma could effectively contribute to the somatic tuning. We selected the spines as a function of distance from the soma (range 100-300 μm; step size 50 μm) and calculated the tuning curve using the same method (Figure S13c, e). When only spines within 150 μm from the soma were selected, the CI did not improve very much (CI = 0.82 within 150 μm) and the results were still far from reproducing the tuning of the soma (Figure S13e). In addition, when only spines above 150 μm from the soma were selected, the CI was smaller than the value of within 150 μm (CI=0.67 above 150 μm). Spines with a similar preferred orientation to the soma showed no apparent distributional bias with distance from the soma (Figure S13d). Taken together, the tuning curve estimated by a simple summation of the number of responding spines was not improved by considering the weighting factors of either the spine signal change or the electrotonic length constant.

### Two forms of functional clustering of similarly tuned spines on the dendritic branch

It has been suggested that, given the impedance matching at the branch point, each dendritic branch may be an electrical compartment for the local processing of input signals and that branch-specific computation may occur (**Branco & Hausser, 2010**). We considered dendritic branches as the integrating unit of synaptic inputs and focused our analysis on dendritic branches (see Discussion). Recent studies have suggested that nonlinear local summation of clustered synaptic inputs on dendrites may influence somatic spiking (**Jia et al., 2010; Wilson et al., 2016; Smith et al., 2013; Major et al., 2013; Palmer et al., 2014; Bicknell & Hausser, 2021**).

Our reconstructed functional input maps (e.g., Figure 3a, c) indicated the clustering of similarly tuned spines in some branches. Therefore, we quantitatively analyzed the spatial location and orientation tuning of spines on each dendritic branch. First, we calculated the distance and the difference in preferred orientation between the spines in each branch (Figure 5). We found that the ΔOri of nearby spines (< 3 μm apart) in the same dendritic branch was smaller in the actual data than in the shuffled data (Figure 5a, 1,000 shuffles of all spines within a segment, p=1 Mann–Whitney U-test with the Bonferroni correction; Figure 5b, 1,000 shuffles of all spines within a cell, p=0.023 Mann–Whitney U-test with the Bonferroni correction). The results indicate the presence of functional clustering of spines at the spatial scale of less than 3 μm distance between spines (neighboring spine-pairs). Next, we counted the number of spines that responded to each orientation stimulus on individual dendritic branches (Figure S14). So far, we found two forms of functional clustering of similarly tuned spines on the dendritic branch, one is clustering of neighbor spine-pairs and the other is the clustering on the dendritic branch (clustered dendrites).

**Figure 5.**
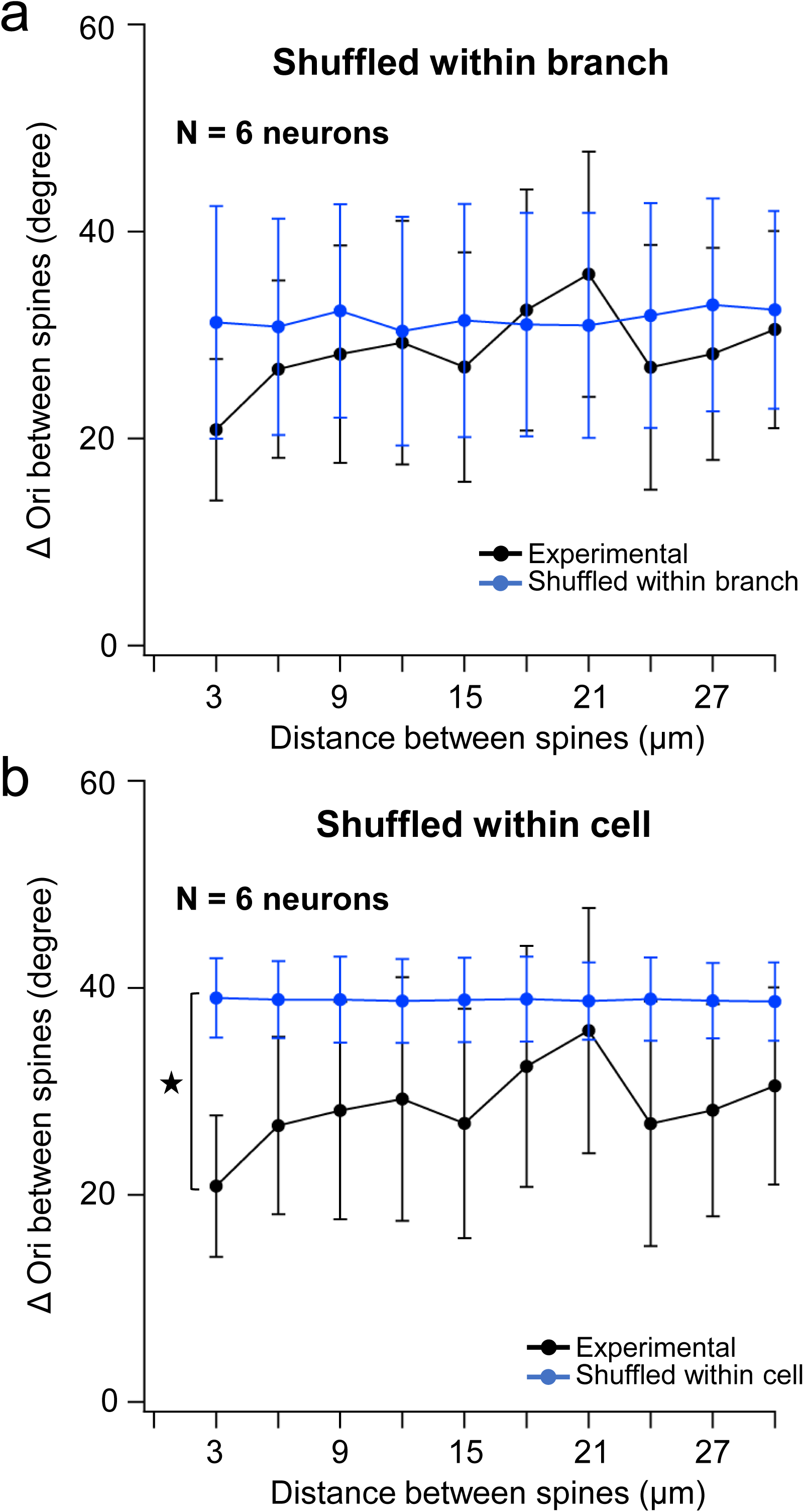
Functional clustering at the spine-pair level. The difference in preferred orientation (ΔOrientation) are plotted as a function of the distance between spines on each dendritic branch and the actual data (**a, b**, black line) are compared with the shuffled data (**a, b**, blue line). In the shuffled data, all the pines were shuffled, while maintaining the original spine locations and the same proportion of orientation-tuned spines. Shuffling was done either across cell (**a**, blue line) or within each dendritic branch (**b**, blue line). ΔOrientation between neighboring spines (< 3μm) of actual data is significantly smaller than that of shuffled data across a cell (**a**, p = 0.023 Mann–Whitney U-test with the Bonferroni correction) and smaller but not significantly different than that shuffled within a branch (**b**, p=0.83 Mann–Whitney U-test with the Bonferroni correction).

We observed clusters of spines with similar responses on a small subset of dendritic branches, but most dendritic branches have either diversely responsive or visually non-responsive spines (Figure S14; see also Figure 3). The proportion of visually responsive spines on each dendritic branch is highly variable, ranging from 0% to 100% in our recorded data (average 20.7% among 208 dendritic branches from 6 neurons). Our functional synaptic input maps (e.g. Figure 3) suggest that the dendritic branches can be classified into three patterns of synaptic input architecture.: branches with clusters of spines of similar responses, branches with spines of diverse responses, and branches with spines where the majority of them show no visual response.

### Relationship between two forms of functional clustering

We examined the relationship between two types of spatial and functional organization of similarly tuned spines: neighboring spine pairs and clustered dendrites. A neighboring spine pair was defined as a pair of similarly tuned spines separated by less than 3 μm, whereas a clustered dendrite was defined as a dendritic branch containing no less than 5–8 similarly tuned spines. We therefore asked whether branch clustering contributes to the occurrence of neighboring spine pairs.

To investigate this relationship, we evaluated the proportion of similarly tuned spine pairs among all spine pairs at different distances along each dendritic branch (30 μm in 3-μm bins) and compared the experimental data with randomized data (Figure S15). For each neuron, 100,000 randomized functional spine input maps were generated. Randomization was performed either within each branch (Figure S15a, blue line) or within the entire cell (Figure S15a, red line). The randomized maps preserved the proportion of visually responsive spines and the original spine positions, while the response properties of the spines were randomly shuffled.

The proportion of similarly tuned spine pairs in the distance bin below 3 μm represents the occurrence of neighboring spine pairs. Only a small proportion of all spine pairs were similarly tuned in the experimental data (Figure S15b, black bar; ∼4.7%, corresponding to ∼44 spine pairs per neuron on average across six neurons). Neighboring spine pairs can arise from the spatial distribution of functional spine inputs within individual branches, branch clustering, and the overall proportion of similarly tuned spine inputs. We analyzed the data by comparing the experimental values with the 95% intervals of the shuffled distributions. The experimental values fell outside the 95% intervals of the shuffled distributions for both comparisons. The corresponding Z statistics were 5.41 for the comparison between “Neighboring spine pairs” and the “Combination of spine input patterns in the dendritic branch,” and 18.04 for the comparison between “Neighboring spine pairs” and the “Combination of branch clustering.” Both Z statistics yielded two-sided p-values of p < 0.001. Thus, based on these randomization analyses, branch clustering may account for approximately half of the excess similarly tuned neighboring spine pairs.

### Predicting somatic tuning from clustering of spines on dendrites and thresholding at soma

We tested a cluster model in which clustering of similarly activated spines can sharpen the broad tuning predicted by a simple summation of activated spines. In this model, we assumed that activated dendritic signals propagate to the soma only when the number of responsive spines in a dendritic branch reaches or exceeds a clustering threshold, and that these spines contribute to somatic depolarization (Figure 6e, f and Figure S16a). As the cluster threshold increased, the proportion of the spines responding to the same preferred orientation as the soma (Figure S16a, red line) among the spines responding to any orientation (Figure 16a, black line) increased (Figure S16a, blue line). Thus, as the threshold increased, the predicted time course became tighter (Figure 6e) and the predicated soma selectivity became sharper (Figure 6f).

**Figure 6.**
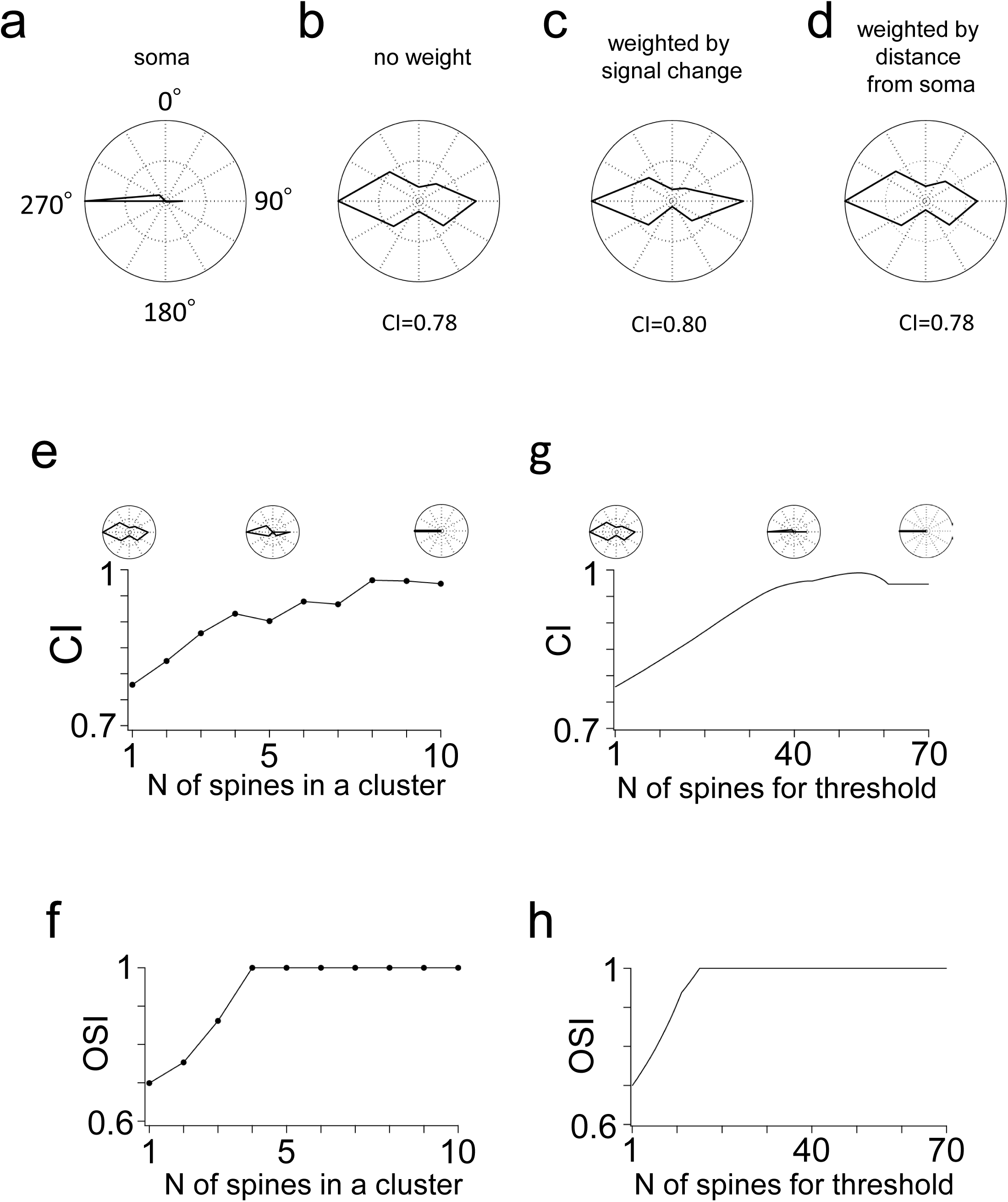
The prediction of somatic tuning was improved by the cluster model on the dendrites and the threshold model at the soma. **a.** A Polar plot of the somatic tuning curve of neuron 5. b. A polar plot from the summation of the number of responsive spines along the entire dendrite. The CI between the soma and the summation of the number of responsive spines increased from the simple summation of spine time courses shown in Figure 4d right. c. The responsive spines are weighted by the individual spine signal changes and then summed. The CI indicates almost no improvement in the prediction of the tuning curve. d. The responsive spines are weighted by the distance from the soma to the individual spines and then summed. The polar plot shows no improvement in the tuning curve. **e.** The CI between the actual and predicted tuning curves of the soma plotted as a function of the number of spines in a cluster. Polar plots of the actual and predicted tuning curves of the soma are shown. **f.** The OSI plotted as a function of the number of spines in a cluster. **g.** The CI between the actual and predicted tuning curves of the soma plotted as a function of the number of spines for the threshold. Polar plots of the predicted tuning curve of the soma are shown. **h**. The OSI plotted as a function of the number of spines used as a threshold.

On the other hand, it has been suggested that whether the soma evokes action potentials or not is controlled by the nonlinear effect of the spike threshold in the soma (**Priebe & Ferster, 2008**). Therefore, we investigated another model to predict the soma tuning. We considered a threshold at the soma (soma threshold) for triggering spikes and investigated whether the prediction of the somatic tuning could be improved when the soma threshold was varied (a threshold model, Figure 6g, h and Figure S16b). In this model, any responsive spine signal propagates to the soma regardless of clustering, and the soma threshold is defined as the number of responsive spines required to trigger a spike in the soma. As the soma threshold increased, the proportion of the spines responding to the preferred orientation of the soma among the spines responding to any orientation increased (Figure S16b). Thus, as the threshold increased, the predicted time course became tighter (Figure 6g) and the predicated soma selectivity became sharper (Figure 6h). These results indicate that both the cluster and soma threshold models are effective models for predicting the soma tuning.

Previous model study suggested the two-layer input-output function of the pyramidal neuron, that is, synaptic input summation at the dendrite occurs in the first layer and thresholded final response at the soma is produced in the second layer (**Poirazi et al., 2003**). This model has been experimentally tested and revealed that combination of dendritic nonlinearity and spike thresholding contribute to the sharpness of orientation tuning in the ferret visual cortex (**Wilson et al., 2016**). Therefore, we tested this model using our experimental data if we see further improvement of tuning also in the mouse (Figure 7). The heat map of CI showed that the highest CI (CI = 1.00) was obtained at the combination of cluster size 8 and soma threshold 7 (Figure 7a, left), and this combination accurately predicted the soma tuning (Figure 7a, right). Another example of a neuron showed a similar improvement in the estimation of the somatic tuning (Figure 7b, CI = 0.99, cluster size 6 and soma threshold 8). A similar trend was observed in the population average of the individual heat maps of the six neurons (Figure 7c). The parameters that best predicted the soma tuning varied from about 6 to 8 spines for the cluster size and from about 5 to 20 spines for the soma threshold (Figure 7c, CI = 0.95, cluster size 6 and soma threshold 6). These results suggest that both local integration in dendrites and global integration at the soma sharpen the broad tuning and determine the fine tuning of the soma (Figure S17).

**Figure 7.**
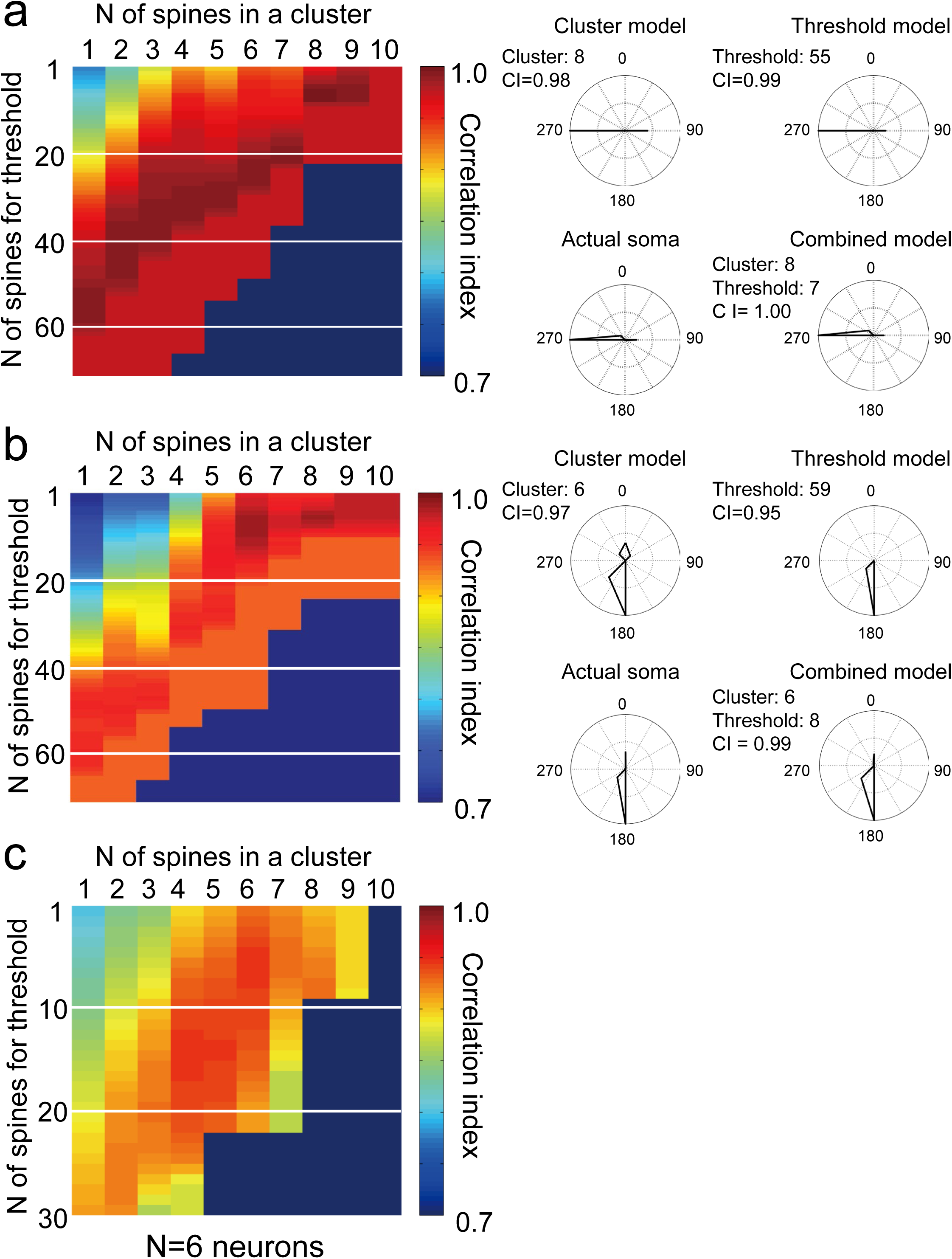
The combined model further improves soma tuning. **a.** A combined model is applied to neuron 5 (see Table 1). CIs are calculated at various combinations of values for the number of spines (1-10 spines) in a cluster and the number of spines (0-70 spines) used as the threshold, and CI values are shown as a heat map. **b.** Another example of a combined model in neuron 2. **c.** Population averaged data from six neurons. Heat maps are calculated for each neuron and averaged.

### Small number of clustered branches and spines are sufficient to reproduce the tuned output signal

Our synaptic input integration model-a cluster model (Figure S16a and Figure 7: cluster model) is able to predict somatic output tuning using only dendritic branches containing clusters of similarly tuned synaptic inputs (5-8 spines/branch), while ignoring other dendrites (Figure 8f). This means that the branches with small number of spines can be considered as functional clusters, suggesting that they are critical branches for soma output tuning. Based on our synaptic input integration model illustrated in Figure S16 and Figure 7, we defined the minimum number of similarly tuned spines as a cluster on a dendritic branch that can reproduce the somatic output tuning. Using this number, we extracted clustered dendrites from the experimental data, reconstructed spatial distribution maps of clustered dendrites for each neuron, and analyzed the proportion of clustered branches as well as the proportion of similarly tuned spines belonging to cluster (clustered spines) in each neuron (Figure 8a-e). The results showed that the number of branches averaged 7 per neuron (Figure 8b; 21.9% of all branches: 12.2%∼31.2% from 6 neurons) and the number of clustered spines averaged 63 per neuron (Figure 8c; 6.9% of all spines: 4.7%∼8.5% from 6 neurons). These dendritic branches are distributed throughout the dendrites without bias (Figure 8a). These dendrites distributed most abundant 50-100 μm from the soma (Figure 9d, 30 of 44 branches in 6 neurons, approximately 68% of the total) and in the second and third branches (Figure 8e, 36 of 44 branches in 6 neurons, approximately 82% of the total).

**Figure 8.**
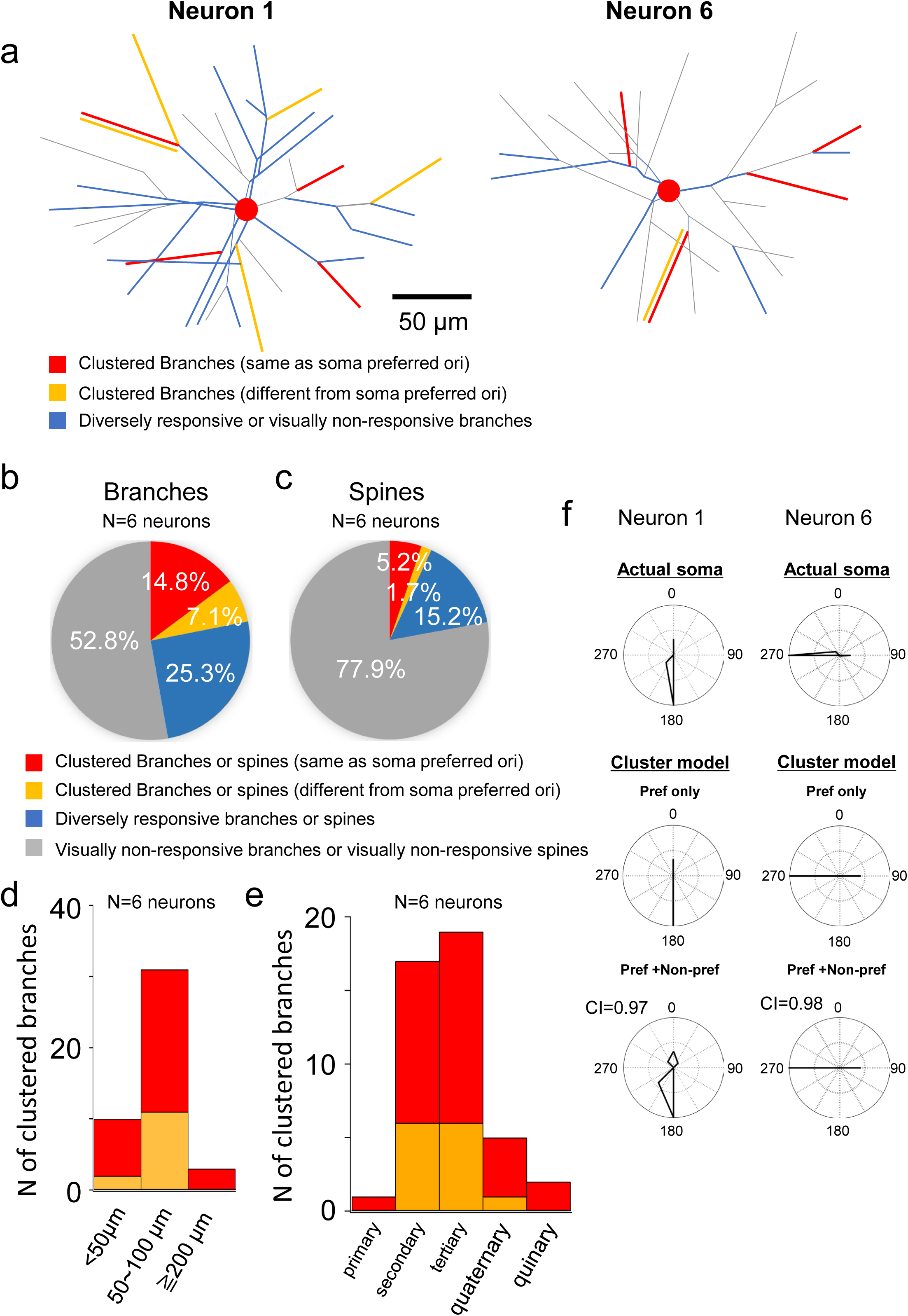
Analysis of dendritic branches with cluster of spines responding to the similar orientations. **a.** Two examples of neuron showing the segments with cluster of spines responding to the same orientation as the soma preferred orientation (red) and different orientation from the soma preferred orientation (yellow). Clustered segments are distributed spatially uniform manner on the dendritic field in both neurons. **b.** Proportion of clustered segments (red and yellow) and non-clustered segments (blue). Among clustered segments (21.9%), the segments of spines responding to the same orientation as the soma preferred orientation (red) is twice as much as the segments of spines responding to the different orientation to the soma (yellow). **c.** Proportion of spines on the clustered (red and yellow) and non-clustered (blue) dendritic segments. Among spines on the clustered segments, the spines responding to the same orientation as the soma preferred orientation (red) is three times as much as the spines responding to the different orientation to the soma preferred orientation (yellow). **d, e.** Distribution profiles of clustered segments on the dendritic field are shown. Number of clustered segments is plotted as a function of the distance from the soma (**d**) and the branch order (**e**). **f.** Prediction of soma tuning using spine responses on the clustered dendrites. Two examples from the same neurons shown in (**a**). upper plot: Actual soma tuning. middle plot: Predicted tuning using spine responses on the clustered segments with the same preferred orientation as the soma. lower plot: Predicted tuning using spine responses on the clustered segments with both the same and the different preferred orientation to the soma. Better prediction was obtained when using the both clustered segments.

We observed that 52.8% (range: 26.8–80.5%, SD = 21.5%) of dendrites had more than 80% visually non-responsive spines; 21.9% (range: 12.2–31.3%, SD = 6.9%) had clusters of spines with similar responses; and 25.3% (range: 6.7–51.2%, SD = 18.4%) had spines with diverse responses (Figure 8b). We assessed whether the proportion of dendrites with predominantly non-responsive spines was greater than expected by chance by performing 100,000 shuffles within individual neurons. In the population (N = 6 neurons), we compared the experimental data (52.8% ± 21.5%, mean ± SD) with the shuffled data (46.8% ± 23.7%, mean ± SD; 100,000 shuffles, N = 6 neurons). The test statistic (Z) was calculated as 0.25, yielding a p-value of 0.802 (two-sided test). This result suggests that the occurrence of dendrites dominated by non-responsive spines can arise from the overall distribution of visually responsive and non-responsive spines, without requiring additional branch-specific organization.

## DISCUSSIONS

In this study, we first established a new method to prevent BAP signal along dendrites, which has been the major obstacle to accurately recording of signal changes in individual spines. Using this method, we conducted large-scale spine imaging (∼1,000 spines per neuron) from layer 2/3 pyramidal neurons in mouse primary visual cortex and obtained new findings. First, functional architecture of dendrites can be classified into three synaptic input patterns.: dendrites with clusters of spines of similar responses, dendrites with spines of diverse responses, and dendrites with spines where the majority of them show no visual response. Next, to predict the sharp output tuning, we examined several theoretical models based on our experimental data and found that the dendritic cluster model, which only considers the dendrites with cluster of spines of similar responses for the control of output signal, is effective in predicting the sharp output tuning. Our data suggest that only a small fraction of spines (6.9% of the total spines) form clusters of similar inputs on a small subset of dendrites (21.9% of total dendrites), and this functional architecture effectively contributes to the generation of sharply tuned output.

### New method to record spine activity by preventing BAP

Imaging of spine activity under the hyperpolarization of somatic membrane potential to prevent BAP has previously been performed by patch-clamping the cell body (**Jia et al., 2010; Chen et al., 2011**) or microinjecting a sodium channel blocker into the soma (**Levy et al., 2012**). In contrast, in this study, we hyperpolarized the somatic membrane potential by using a long-lasting bistable inhibitory opsin (**Berndt et al., 2016**) with a soma-targeting signal. Our optogenetic method is technically easier and can maintain the membrane potential longer than electrophysiological or chemical methods to perform large-scale dendritic mapping. On the other hand, there is concern that a continuous increase in chloride conductance due to activation of soma SwiChR++ channels may affect synaptic inputs. Increased chloride ions in the soma may to some extent spread to dendrites. Normally, intracellular chloride levels are regulated by the neuron-specific potassium chloride transporter (KCC2) expressed in both the soma and dendrites (**Baldi et al. 2010**). However, it is unclear how effectively dendritic KCC2 prevent the spread of chloride ions from the soma to the dendrites in our experimental condition. The lack of difference in the magnitude of synaptic responses before and after photoinhibition at non-preferred orientations (Figure 2C), which are thought to be largely unaffected by BAP, suggests that the increased chloride ion permeability of the soma has not significant effect on the dendrites.

### Large-scale spine imaging

To understand the tuning mechanisms from synaptic input to soma output, it is necessary to record spine activity from individual neurons as much as possible. It has been previously reported that a single layer 2/3 pyramidal neuron in the cortex possess ∼7,000 spines and among them ∼4,000 spines are located on basal dendrites (**Lascone et al., 2020**). In this study, we performed large-scale functional imaging of spines on the basal dendrites. We recorded ∼1,000 spine activities from each neuron in response to visual stimulation. We found that on average only 21.6% of the spines are visually responsive to the drifting grating stimulation. It is unexpectedly small that most of the synaptic inputs to layer 2/3 pyramidal neurons in V1 are visually unresponsive. The proportion of visually unresponsive spines was not related to the time course of the recording. For example, in Figure S14, the higher the dendrite number, the later they were recorded from the start, but there is no relationship between the dendrite number and the presence or absence of a visual response. Our results showed that the proportion of visually responsive spines on each dendritic branch is highly variable, ranging from 0% to 100% in our recorded data (average 20.7% among 208 dendritic branches from 6 neurons). This suggests that the synaptic input organization of each dendritic branch is functionally heterogeneous. A small subset of dendritic branches has clusters of spines with similar orientation tuning, while others have spines with heterogeneous orientation tuning or no visual response. The origin and synaptic organization of visually unresponsive inputs to a single neuron is not known well. One of the possible origins could be visually unresponsive surrounding neurons within V1. Previous studies by ours and others have shown that only about half of the excitatory neurons in V1 are visually responsive to the drifting grating stimulus at a given spatial and temporal frequency (about 50-60% in layer 2/3 of V1, **Kerlin et al., 2010; Marshel et al., 2011; Kondo et al., 2016**). Recurrent connections are mostly formed on the basal dendrites (**Spruston, 2008**), from which we recorded spine signals in this study. Another possible source of visually unresponsive inputs may be projections from outside V1. Recent retrograde labeling studies have shown that CTB (retrograde labeling tracer) injections into V1 layer 2/3 revealed inputs from many nonvisual brain areas (**Morimoto et al., 2021**). In summary, a single layer 2/3 pyramidal neuron in V1 may receive mixed synaptic inputs from both visually responsive and visually unresponsive neurons inside and outside of V1, and the dendritic branches may be organized by the spines that receive visually responsive and visually unresponsive synaptic inputs according to a rule that remains to be clarified.

### Functional clustering of synaptic inputs on subsets of dendritic branches

We found that the spines evoked by the preferred soma orientation were dispersed across multiple dendritic branches, but clustered on some branches of layer 2/3 pyramidal neurons in mouse V1. Our simulated model suggested that as the number of spines used to define a cluster increased, the predictive accuracy of somatic tuning increased (Figure 7b). Large-scale spine imaging allowed us to estimate the number and the spatial distribution of clustered dendritic branches and the number of spines in the cluster. We found that only a small fraction of dendritic branches (∼22% of the total number of dendrites) that are spatially distributed can reproduce somatic tuning without considering other dendritic branches. Accordingly, the number of spines clustered on dendritic branches is also small (∼7% of the total number of synaptic inputs). This type of functional organization may be conserved across species with or without functional columns and different visual functions, such as functional clustering for spatial location in mice (**Iacaruso et al., 2017**), clustering of callosal and non-callosal inputs in mice (**Lee et al., 2019**), and functional clustering of orientation inputs in ferrets (**Wilson et al., 2016**).Functional clustering of synaptic inputs may be effective for strong activation while maintaining robustness against information loss along dendrites (**Stuart and Spruston, 2015**).

### Weighted and unweighted summation of activated spines failed to estimate the sharp tuning of the soma

Previous studies (**Chen et al., 2013; Wilson et al., 2016**) and our study showed that the preferred orientation of the soma can be predicted by the summed number of activated spines or the averaged time course of spine responses, but showed broader orientation tuning. Even when we considered the magnitude of the spine signal change or the distance between the soma and the spine, the estimation did not improve. Regarding the magnitude of spine signal change, calcium signaling is not the direct measure of synaptic transmission via AMPA receptors, but rather via NMDA receptors (**Sobczyk et al., 2016**). Recent work (**Scholl et al., 2021**) suggests that the magnitude of calcium signal change may not be a good predictor of synaptic strength. Regarding the distance between the soma and activated spines, it has been previously shown the distance independence of synaptic inputs on the somatic synaptic integration (**Magee, 2000**). Thus, our result is not surprising given the long length constant (∼300 μm) of cortical layer 2/3 pyramidal neurons (**Larkum et al., 2007**) and the distance-independent distribution of activated spines with different orientation selectivity (Figure S14d).

### Cluster model at the dendrites to predict soma tuning

The spatial arrangement of functional synaptic inputs on the dendrite should provide useful information to reveal the computation performed by dendritic integration. Many theoretical models have been developed to understand the electrical behavior of dendrites (**Poirazi & Papoutsi, 2020**). In the compartmental model, dendrites are divided into multiple compartments, and each compartment is considered as a computational unit (**Rall, 1964; Poirazi et al., 2003)**. Dendrites can be divided into compartments in various ways. Among them, considering the impedance matching at the branch point, each dendritic branch can be an electrical compartment for the local processing of input signals, and branch-specific computation could occur (**Branco & Hausser, 2010**). The electrically isopotential dendritic length at which synaptic inputs can be effectively integrated, is reported to be ∼40 μm in neocortical pyramidal neurons (**Mel, 1993**). Because the average length of the dendritic branches in our recorded neurons was mostly around 20∼30 μm (see Methods), we considered the dendritic branches as the integrating unit of synaptic inputs and focused our analysis on the dendritic branches. We observed clusters of similarly tuned synaptic inputs on the dendritic branches of layer 2/3 pyramidal neurons. This feature is particularly efficient for the somatic spiking because unlike spatially distributed coactivated spines, clustered coactivated spines can evoke the dendritic spikes and influence the somatic activity more effectively (**Stuart & Spruston, 2015; Ujfalussy & Makara, 2020**). Dendritic spikes triggered by coactive synaptic inputs have been observed as dendritic hotspots in the previous studies (**Jia et al., 2010; Wilson et al., 2016**). In our present study, the cluster model improved the estimation of the soma orientation tuning, suggesting the importance of the clustering of synaptic inputs with similar orientation tuning. The dendritic hotspots were not observed in our recording, probably because the strong hyperpolarization of the soma with the optogenetic method may reduce the probability of triggering the dendritic spikes without significantly affecting the spine signals. Note that, the somatic tuning was measured before photoinhibition of the soma activity, so the measurement of the somatic tuning was not affected by photoinhibition.

### Threshold model at the soma to predict somatic tuning

Previous studies (**Chen et al., 2013; Wilson et al., 2016; Scholl et al., 2021**) and our study showed that the output tuning was narrower than the total input tuning predicted by simple summation of the inputs. It has also been reported that the tuning curve from the membrane potential is broader than that from soma spiking (**Troyer et al., 1998; Finn et al., 2007; Lien & Scanziani, 2013**). To explain the sharp tuning of the soma in V1 neurons, a spike-threshold model has been proposed in which only the preferred orientation raises the membrane potential above the threshold and triggers an action potential in the soma (**Priebe & Ferster, 2008**). In our model, we introduced an analogous spike-threshold model for V1 neurons. We assumed that the number of active spines corresponds to the amplitude of the excitatory postsynaptic potential (EPSP) that propagates to the soma and contributes to the level of the soma membrane potential level. We hypothesized that the somatic action potential is evoked only when the number of active spines is above a certain threshold. By applying the threshold model, the tuning of the somatic activity to visual stimulation was sharpened and more accurately predicted, suggesting the importance of thresholding for the output tuning at the soma.

### The combined model is better for predicting somatic tuning from synaptic inputs

Previous model study suggested the two-layer input-output function of the pyramidal neuron, that is, synaptic input summation at the dendrite occurs in the first layer and thresholded final response at the soma is produced in the second layer (**Poirazi et al., 2003**). This model has been experimentally tested and revealed that combination of dendritic nonlinearity and spike thresholding contribute to the sharpness of orientation tuning in the ferret visual cortex (**Wilson et al., 2016**). Therefore, we tested this model using our experimental data if we see further improvement of tuning also in the mice.

The combination of cluster and threshold models improved the prediction of the somatic tuning better than either the individual model. Two steps of integration finally sculps to the sharply tuned output signal. By dividing information processing into two compartments, the soma and the dendrites, neurons may acquire a greater computational capacity than either the cluster or threshold model of integration. Our data are consistent with the previous findings in ferrets (**Wilson et al., 2016**) and this two-step sharpening of the output signal may be a common mechanism across species.

### Effects of inhibitory inputs on soma output tuning

Excitatory neurons in V1 not only receive excitatory inputs from other excitatory neurons, but also receive inhibitory inputs from interneurons. One of the potential roles of inhibitory inputs for the orientation tuning is to sharpen the soma output of excitatory neurons (**Priebe and Ferster, 2008; Liu et al., 2011; Hansel and Vreeswijk, 2012**). The tuning of inhibitory neurons has been shown to be mostly broad for orientation (**Sohya et al., 2007; Niell and Stryker, 2008; Kerlin et al., 2010**). Such broad hyperpolarizing potentials can raise the overall excitatory threshold for the soma spiking activity and unselectively suppress the responses at all orientations. Hence, in our model, inhibitory effects are equivalent to increasing the threshold and taken into account in the threshold model.

### Role of apical and basal dendritic inputs

Apical and basal dendrites receive inputs from different sources and differ in the density of some types of ion channels (**Stuart & Spruston, 2015**), suggesting their different roles in information processing (**Spruston, 2008**). In the present study, we recorded only the basal dendrites of layer 2/3 neurons in V1, but previous studies showed that the clustering of spines with similar orientation preference was not significantly different between apical and basal dendrites in ferret V1 (**Wilson et al., 2016**) and the removal of apical dendrite does not alter orientation-tuning (**Park et al., 2019**). On the other hand, a close relationship between soma firing and activity in the basal dendrite, but not in the apical tuft has been reported (**Hill et al., 2013**). Indeed, in our study, soma tuning was estimated only from synaptic inputs on basal dendrites. Regarding the role of apical dendrites, several *in vivo* studies have suggested that synaptic inputs to apical dendrites may play an important role in synaptic plasticity (**Cichon & Gan, 2015**), modulation of sensory information (**Smith et al., 2013; Palmer et al., 2014**) or reception of surround information in the visual field (**Roth et al., 2016**).

### Future applications

To understand how neural circuits work, it is important to understand not only the cell types and their connectivity in neural circuits, but also the cell type-specific computation. Cortical neurons can be divided into two types: excitatory and inhibitory neurons. In inhibitory neurons, similar to the excitatory neurons, action potentials initiated in the soma also propagate back into the dendrites (**Kaiser et al., 2001**). Because inhibitory neurons generally lack spines and excitatory inputs are formed directly on the shaft (**Harris & Shepherd, 2015**), calcium signals derived from BAP are completely mixed with the signals of the shaft synapse, the previous regression and subtraction method (**Chen et al., 2013; Wilson et al., 2016; Iacaruso et al., 2017; Scholl et al., 2021**) would be difficult to apply to this type of synaptic input. To our knowledge, only one study has reported the *in vivo* recording of synaptic inputs from inhibitory neurons whose cell bodies do not fire to the visual stimulation (**Chen et al., 2013)**, and synaptic imaging of inhibitory neurons whose cell bodies do fire has not been attempted because of this difficulty. Therefore, our novel method for recording synaptic activity without interference from BAP signals will be particularly useful for the synaptic imaging of inhibitory neurons.

Previous methods for suppressing BAP (**Jia et al., 2010; Chen et al., 2011; Levy et al., 2012**) have not been applicable for repeatable imaging over days; our new method is the first to allow easy and repeatable imaging of synaptic inputs over days *in vivo*. Thus, this new method will contribute to the broader field of synaptic plasticity, which studies studying the dynamic changes in location and activity of synaptic inputs, such as pruning of synaptic inputs during development, pathological alterations of synaptic inputs in the psychiatric disorders and the reorganization of synaptic inputs during learning.

## MATERIALS AND METHODS

### Animals

C57BL/6 wild-type mice or Thy1-GCaMP6s transgenic mice (GP4.3) (2-3 months old, male) were used for all experiments. C57BL/6 mice were purchased from SLC (Hamamatsu, Japan). Thy1-GCaMP6s mice (stock #024275) were purchased from Jackson Laboratory (Bar Harbor, ME, USA). All mice were maintained in the animal facility at the University of Tokyo and housed 2-3 mice per cage in a temperature-controlled animal room with a 12 h/12 h light/dark cycle. All procedures were conducted in accordance with the protocols approved by the University of Tokyo Animal Care and Use Committee (approval number: H22-104, H22-111).

### Local and sparse expression of GCaMP6s and SwiChR++ in V1 layer 2/3 neurons

The genetically encoded calcium indicator GCaMP6s and optogenetic protein SwiChR++ were locally and sparsely expressed using the AAV infection method. A mixture of AAV2/1-hSyn-Cre and AAV2/1-hSyn-flex-GCaMP6s was used for sparse GCaMP6s expression (∼1 × 10^8^ and ∼1 × 10^13^ genome copies/mL, respectively, purchased from the addgene) and AAV2/1-CaMKIIa-mCherry-P2A-Kv2.1-SwiChR++ for SwiChR++ expression (∼1 × 10^13^ genome copies/mL, originally reconstructed from the AAV vector purchased from the Vector Core of Stanford University). The reconstructed vector for SwiChR++ expression contains mCherry to label SwiChR++-expressing neurons and a Kv2.1 fragment to localize SwiChR++ in the soma to prevent excess hyperpolarization of dendrites. Mice were anesthetized with isoflurane (1.5% vaporized in air) and fixed on stereotaxic frames. The skin was incised at the midline, and the periosteum was removed from the skull. A small craniotomy was performed just above V1 at selected stereotaxic coordinates (−3.5 mm from the bregma, 3.0 mm lateral to the midline), and a glass pipette filled with AAV was inserted (0.4 mm depth from the pia). AAV was injected under pressure (500 nL using a NanoJect III, Drummond Scientific, Broomall, PA, USA). The infection area is typically defined as a 400-700 μm diameter from the injection site. Imaging was performed two weeks after AAV infection.

### Imaging

For the imaging of spines and soma of V1 neurons, a craniotomy (4 mm diameter) was performed over V1. The dura mater was removed, and a cranial window was constructed by sealing the opening with a cover slip (No.1, 6.5 mm diameter, Matsunami, Japan). Mice were anesthetized with isoflurane (1.0–2.0%) during surgery. Two-photon calcium imaging of spines and soma was performed under light anesthesia (0.2% isoflurane under sedation with 2.5 mg/kg chlorprothixene). For the spine imaging, neurons that expressed both GCaMP6s and mCherry were selected for imaging. First, imaging of the soma was performed to record the somatic tuning before photoinhibition of somatic activity, and then somatic activity was suppressed by light stimulation and spine imaging was performed. In experiments to examine the effects of light suppression on surrounding cells or spines before and after light suppression, the activity of the cells or spines was first imaged before photoinhibition and then a target cell activity was suppressed by light stimulation and the activity of the same cells or spines was imaged and compared. Imaging was conducted using a two-photon microscope (FVMPE-RS, Olympus, Tokyo, Japan) with a 25× objective lens (XPLN25XWMP2, NA = 1.05, Olympus, Tokyo, Japan) at a wavelength of 920 nm (InSight DS, Spectra Physics, Milpitas, CA, USA). Images were obtained in a 2-D plane at 1 Hz (Galvo scanning; for BAP inhibited method) or 15 Hz (resonant scanning; for BAP subtracted method) (**Chen et al., 2013**). The image size was 512 × 512 pixels (the resolution was 0.125 μm/pixel for spines and 0.5 μm/pixel or 1.0 μm/pixel for soma). The average laser power directed at the sample was modulated between 10 and 40 mW, depending on the imaging depth. To correct the refractive index mismatch in the brain tissue, we carefully adjusted the axial position of the objective lens to obtain the maximum signal intensity.

Visual stimulations were presented on a 32-inch LCD display (ME32B, Samsung) using PsychoPy2 (**Peirce, 2008**). Orientation selectivity was investigated by square-wave drifting gratings in eight directions (45° apart, spatial frequency = 0.04 cpd and temporal frequency = 2 Hz). These eight patterns were presented for ∼4 s each (4 frames) interspersed with blank (uniform) gray stimuli of either the same duration or twice the duration. The stimuli were presented 10 times.

### Optogenetic inhibition of soma activity

Optogenetic inhibition of soma activity was performed by directing a focused visible laser (wavelength: 458 nm, beam diameter: 0.53 μm) through a small round region of interest (ROI) on the soma through an objective lens by Galvano mirror raster scanning (scan size: 20 x 20 pixels (0.5 μm/pixel), scan speed: 100 ms/frame). The laser power was 50 μW after the objective lens. The duration of the laser stimulation was 100 s. The photoinhibition of somatic activity typically lasted for ∼60 min, and somatic activity was repeatedly inhibited before recovery from inhibition during the recording of spine activity.

### Data analysis

All analyses were performed using custom-written programs in MATLAB (MathWorks, Natick, MA, USA). The data was processed between experimental groups in the same manner using the same computer code; thus, randomization and blinding were not necessary for data analysis. We did not perform experiments blindly. To obtain an orientation map, calcium signal changes were calculated for all pixels, and each pixel was colored according to the response to the orientation stimulation [hue: preferred orientation; lightness: response magnitude; saturation: OSI].

The spines and soma were detected manually using a template-matching algorithm with convolution mask images. The centroids of the spines and soma were determined from a mask image of these structures. Time courses of fluorescent change were extracted by averaging a circle around the centroid (spines, 0.18 µm radius; soma, 2.5 µm radius).

To estimate the out-of-focus signal around individual spines or soma, we created ring-shaped masks with pixels within 10 pixels (spines, 1.25 µm; soma, 5 µm) from their edge, excluding pixels less than 4 pixels (spines, 0.5 µm; soma, 2 µm) from the edge of the spines or soma. The out-of-focus signal obtained from these masks was subtracted from the fluorescence signal of the spines and soma (contamination ratio = 0.3) (**Kerlin et al., 2010**).

The orientation selectivity was calculated from the corrected time courses. Visually evoked fluorescent changes were calculated as the change in fluorescence normalized to the baseline fluorescence (dF/F). Baseline fluorescence was obtained from the average of the last two frames during the blank periods. The p value for responsiveness p(resp) was obtained from ANOVA across the blank and stimulus periods, and the p value for selectivity was obtained from ANOVA across stimulus periods.

The preferred orientation was calculated using vector averaging (**Swindale, 1998**), defined by the following equations: a = ΣR_i_ × cos(2θ_i_); b = ΣR_i_ × sin(2θ_i_), θ_pref_ = 0.5arctan(b/a). R_i_ is the response to the i^th^ direction θ_i_ (eight directions, 45°apart, spanning 0−315 degrees). θ_pref_ is the preferred orientation. The orientation selectivity index (OSI) was calculated using the following formula: (R_pref −_ R_ortho_) / (R_pref_ + R_ortho_), where R_pref_ is the response to the preferred orientation and R_ortho_ is the response to the orientation orthogonal to the preferred orientation. Spines or neurons were considered responsive when they met the following criteria: the p value for responsiveness was less than 0.01, and the maximum response was more than 3%. Among responsive spines or neurons, sharply orientation-selective spines were defined when they met the following criteria: the p value for selectiveness was less than 0.01, and the OSI was more than 0.3.

The preferred direction was calculated using a Gaussian fitting method. A tuning curve was fitted with a sum of two circular Gaussian functions (von Mises distribution), and the peak of the fitting curve was considered the preferred direction. As an index of direction selectivity, DSI was calculated using the following formula: DSI = (R_pref −_ R_opp_) / (R_pref_ + R_opp_), where R_pref_ is the response to the preferred direction and R_opp_ is the response to the opposite of the preferred direction. The responsive boutons or neurons were defined using the same criteria as for the orientation analysis. Among responsive boutons or neurons, sharply direction-selective boutons were defined as those that met the following criteria: the p value for selectiveness was less than 0.001, and the DSI was more than 0.3.

Spine activities were calculated by subtracting the scaled dendritic signal, as previously reported^10^. Briefly, dendritic shafts (∼20 µm in length) excluding spines were extracted and ΔF/F (ΔF/F_dendrite) was calculated. Individual spine signals (ΔF/F_spine) were plotted against ΔF/F_dendrite, revealing two components: a dendrite signal component and a spine signal component. To remove the bAP-related dendrite signal component from the spine signals, the scale factor for the dendrite signal component was determined, and the spine-specific signal was calculated as follows: ΔF/F_spine_specific = ΔF/F_spine − α ΔF/F_dendrite. α was determined using robust regression (MATLAB function ‘robustfit.m’) of ΔF/F_spine vs. ΔF/F_dendrite (the slope of the fitted line in Figure S10).

To compare the experimental results with those expected by chance, we generated shuffled maps of functional spine input for each experiment. These randomized maps preserved the original spine positions and the same proportion of orientation-tuned spines, while randomly reassigning the orientation preferences of the spines.

The correlation index (CI) between the actual and estimated tuning curve of the soma was calculated as follows:

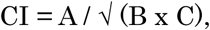

A. = ∑ (actual tuning curve-mean (actual tuning curve)) x (estimated tuning curve-mean (estimated tuning curve))
B. = ∑ (actual tuning curve-mean (actual tuning curve))^2^,
C. = ∑ (estimated tuning curve-mean (estimated tuning curve))^2^.

### Estimation of tuning curve of soma activity by weighting

First, we divided the spines into two types: one was visually responsive (spine activity = 1) and the other was visually unresponsive (spine activity = 0). The effect of visually responding spines on soma tuning was weighted by multiplying either signal change (ΔF/F) or distance between the soma and the given spine (x) by the spine activity value (either 0 or 1). When using signal change as a weighting factor, we estimated the tuning curve of the soma activity as follows:

Tuning curve of soma activity = ∑ (ΔF/F) × (spine activity)

(ΔF/F) is the signal change of the spine to visual stimulation in one of 8 directions. Spine activity at one of 8 directions is assigned 0 or 1 (0: not responsive, 1: responsive). When using the distance between the soma and a given spine as a weighting factor, we estimated the tuning curve of soma activity as follows:

Tuning curve of soma activity = ∑ (exp (− x/L)) × (spine activity)

x is the distance between the soma and a given spine and L is the length constant. L was fixed at 200 µm in Figure 6e and varied between 100 µm, 200 µm, and ∞ in Figure S14a and 13b.

Spine activity at one of 8 directions is assigned 0 or 1 (0: not responsive, 1: responsive).

### Reconstruction of functional synaptic input map

To perform quantitative analyses of the functional geometric organization and synaptic integration model, we reconstructed the functional synaptic input map. Before starting the functional imaging of spines, we took 3-D images of recorded neurons (0.5 µm/pixel) with a 1 µm z-step and reconstructed the anatomical 3-D image. The locations of spines in the functional 2-D images were reconstructed on anatomical 3-D images by aligning the functional 2-D images with the anatomical 3-D image. The coordinates of the spines were defined by setting the coordinates of the center gravity of soma as (0,0,0) and the reconstructed 3-D functional synaptic input map. The distance between the soma and a given spine or between two given spines was calculated as the Euclidean distance using the 3-D functional synaptic input map. The average length of the dendritic branches in our recorded neurons was 24.3 ± 1.1 μm (6 neurons, 214 branches).

### Correlation analysis between two spine activities

To calculate the correlation of the two spine activities, we applied the shuffle-corrected cross-correlation method. First, we calculated trial-averaged time-course data for each stimulus as the shuffle-predictor of visual response. The shuffle-corrected time-course data were obtained by subtracting the shuffle-predictor from the original time-course data for each stimulus. The shuffle-corrected cross-correlogram was obtained as a cross-correlogram between shuffle-corrected time courses. This procedure is equivalent to repeating the shuffle correction for every possible shuffle. The correlation coefficients (values of the shuffle-corrected cross-correlogram at time zero) were calculated between the two spines.

### Analysis for the relationship between two forms of functional clustering

We defined “clustering” as two types: one is the ‘neighboring spine-pairs’, and the other is the ‘clustered dendrites. Neighboring spine-pairs refers to clusters of similarly responding spines located within 3 μm of each other. Clustered dendrites refer to clusters of similarly responding spines (5-8 spines within a dendritic branch exhibited similar responses) situated on the same dendritic branch. In neighboring spine-pairs, the distance between similarly responding spines is less than 3 μm, regardless of whether the spines are located on opposite or the same sides of the dendrite.

To analyze the relationship between the clustering at the neighboring spine-pairs and at the clustered dendrites, we calculated the correlation-coefficient between the number of similarly responded spine pairs and the number of similarly responded spines on each dendritic branch. Then, we generated 100,000 new synaptic input maps (randomized maps) for each experiment. Randomization was done either within each dendritic branch or within a cell. These randomized maps preserved the 3-D original spine positions and the same proportion of orientation-tuned spines, while randomly reassigning the orientation preferences of the spines. From these randomly reassigned maps, we calculated the correlation-coefficient as described above and statistically compared the correlation-coefficient values from the experimental map and two different randomized maps.

### Cluster model analysis

First, we counted the number of visually responding spines in each direction on individual dendritic branches. We set the variable threshold (1–10) for the number of spines in each direction of visual stimulation. If the number of spines exceeded the threshold, we regarded these spines as contributors to soma activity. We repeated this analysis on all the dendritic branches and summed the number of responding spines for each direction of visual stimulation. Finally, we estimated the tuning curve of soma activity from the summed number of spines in each direction of visual stimulation.

### Threshold model analysis

In the threshold model, we regarded all the visually responsive spines as contributors to the soma activity, but somatic spikes occurred only when the number of summed responding spines at the soma was above the threshold number of spines. We varied the threshold, and the tuning curve of soma activity was estimated by subtracting the threshold number of spines from the summed number of spines.

### Classification of dendrites

To classify dendrites as “visually responsive” or “visually non-responsive,” we defined dendrites in which more than 20% of the spines were visually responsive was classified as a “visually responsive dendrite.” The remaining dendrites were classified as “visually non-responsive”. Visually responsive dendrites were further divided into two groups: dendrites containing clusters of spines with similar visual responses (“clustered dendrites”) and dendrites containing spines with diverse visual responses (“diverse dendrites”). We defined a “clustered dendrite” as a dendritic branch containing no less than 5-8 similarly tuned spines. The number of spines used to define a cluster varied among neurons and was determined based on the number of spines that yielded the highest correlation coefficient between the experimentally measured somatic tuning curve and the somatic tuning curve predicted by the model, as shown in Figure 7.

### Immunohistochemistry

AAV infected mouse (two weeks after infection) was fixed by perfusing 4 % paraformaldehyde solution via vasculature system through heart. Brain was removed and post-fixed in the same fixative solution for overnight at 4 degrees. Postfixed brain was cryopreserved by dehydration with sucrose solution, starting at 10% concentration and gradually increasing to 20% and 30%. Coronal slice (50 μm thickness) from primary visual cortex was prepared by a cryostat (CM1520, Leica). Brain sections were run through an immunohistochemistry protocol in order to reveal the hemagglutinin (HA) tag fused to the SwiChR++ as follows. (1) blocking 1h with 5% goat serum + 2.5% Bovine Serum Albumin + 0.1% Triton X-100 in 0.01M PBS; (2) reacted with primary antibody (High affinity anti-HA rat monoclonal antibody (clone3F10), catalogue# 11867423001, Roche), 1:200, 24h at 4 C; (3) washed with PBS at room temperature, 3 times, 15 min for each; (4) reacted with secondary antibody tagged with Alexa Fluor 488 (goat anti-rat IgG, product number A11006, Invitrogen), 1:200, 2-3h at room temperature; (5) washed with PBS at room temperature, 3 times, 15 min for each. Brain sections were mounted with aqueous mounting solution (ProLong Gold, Thermofisher) and analyzed by confocal microscopy (A1R, Nikon).

### Statistical analysis

All data are presented as the mean ± standard error of the mean (SEM) unless stated otherwise. No statistical methods were used to pre-determine sample sizes, but our sample sizes are similar to those generally employed in the field. A one-sample t-test was used to compare a single sample mean to a null hypothesis value. A paired-sample t-test was used to compare between paired observations. A two-sided Mann-Whitney U-test or an unpaired-sample t-test was used to compare two independent groups. An ANOVA and Bonferroni-corrected post hoc test were performed when more than two groups were compared. For analyses involving shuffled data, a null distribution was generated by repeatedly shuffling the data within each cell. The 95% interval of the shuffled distribution was calculated, and the Z-score was calculated as (observed value − mean of the shuffled distribution) divided by the standard deviation of the shuffled distribution. Two-sided P values were then calculated from the corresponding Z-scores.

## ACKNOWLEDGEMENTS

We thank Dr. M. Uemura and Virus Vector Core (Prof. Kobayashi’s Group) of Brain/MINDS (Brain Mapping by Integrated Neurotechnologies for Disease Studies) for the production of AAV; A. Hayashi, Y. Kato, M. Taki, T. Inoue, Y. Sono, A. Honda for animal care and genotyping; N. Yamamoto and T. Ohmine for lab administration; all of the members of Ohki laboratory for support and discussion; The Genetically-Encoded Neuronal Indicator and Effector (GENIE) Project for providing GCaMP6s. Prof. K. Deisseroth for providing SwiChR++.

This work was supported by Brain/MINDS and Brain/MINDS2.0 from AMED (14533320, JP16dm0207034, JP20dm0207048, JP21dm0207014, JP23wm0625001, JP24wm0625203 to K.O., JP21dm0207111 to Dr. H. Hirai and the Brain/MINDS AAV vector core); Institute for AI and Beyond (to K.O.); CREST-JST (JPMJCR22P1 to K.O.), JSPS KAKENHI (Grant numbers 17H03540, 20H03336, 21H05165, 23H02572, 24H02331 to S.K., 25221001, 19H05642, 20H05917 to K.O.), AMED (Grant number JP21wm0525013 to S.K.), The Mitsubishi Foundation to S.K…

## AUTHOR CONTRIBUTIONS

S.K. and K.O. designed the research. S.K. and K.K. performed the experiments and analyzed the data. S.K. and K.O. wrote the manuscript.

## DATA AND CODE AVAILABILITY

The data that support the findings of this study and all custom code used in this study are available from the corresponding author upon reasonable request.

---

**Figure S1.**
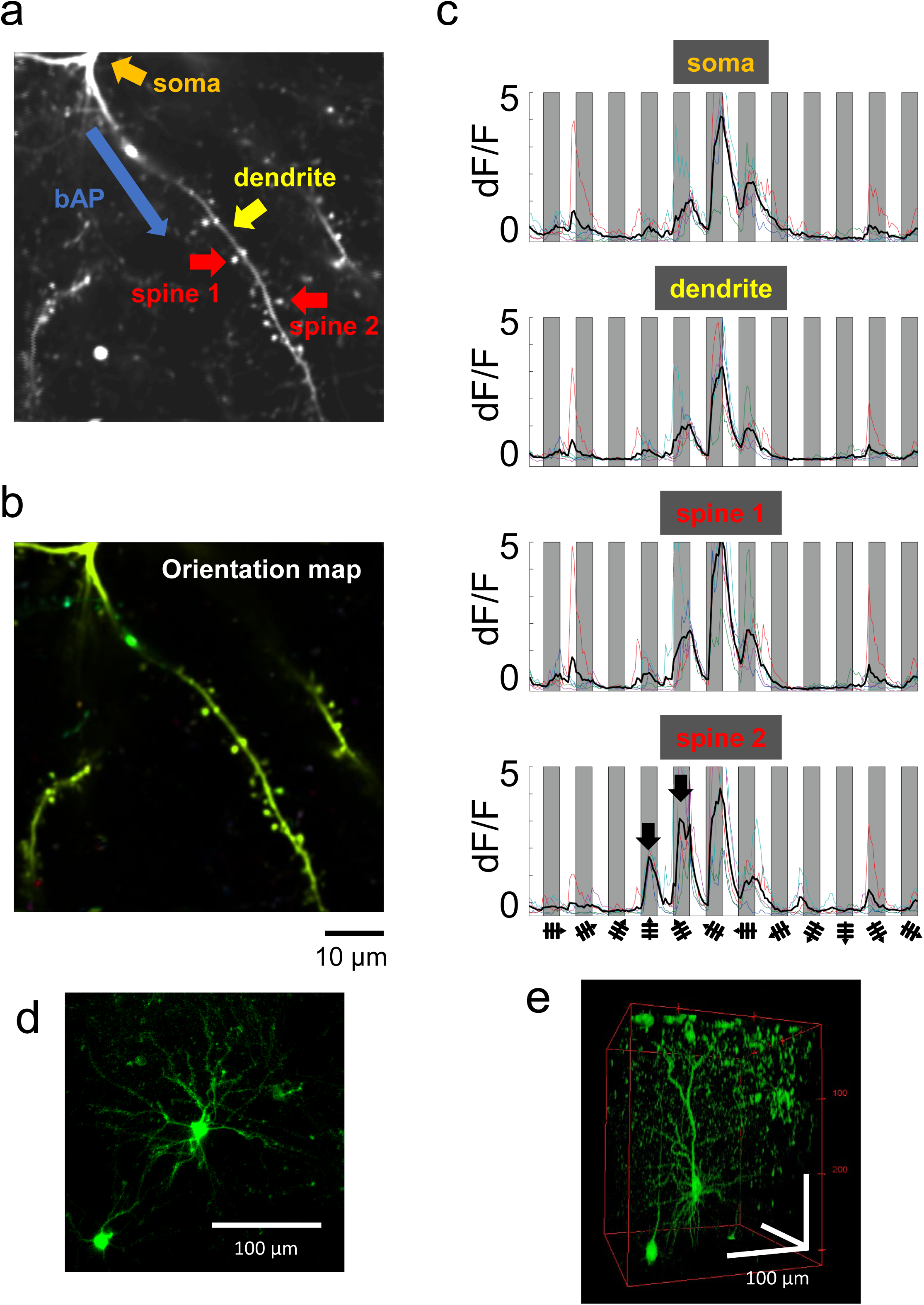
BAP signals along dendrites interfere with spine activity. **a.** An example frame of recording of BAPs originates from the soma and travel backward through the dendritic arbors, where they interfere with spine signals. **b.** The orientation map is shown, and the dendrites and spines are labeled with the same color (green, horizontal orientation stimulation). **c.** Calcium signal changes to the visual stimulation are shown for the soma, dendrite and two spines. The lines with different colors are the responses to each single trial (10 trials in total), and the black lines are the averaged time course of 10 trials. The time courses of these elements show similar shapes, indicating that spine signals are masked by BAP signals. Note: in spine 2, the signal changes indicated by black arrows are different from those of the dendrite. These signals may include original spine signals. d, e. Sparse expression of GCaMP6s in our experiments. Two-dimensional image perpendicular to the cortical surface (d). Three-dimensional image reconstructed from a stack of two-dimensional images (e).

**Figure S2.**
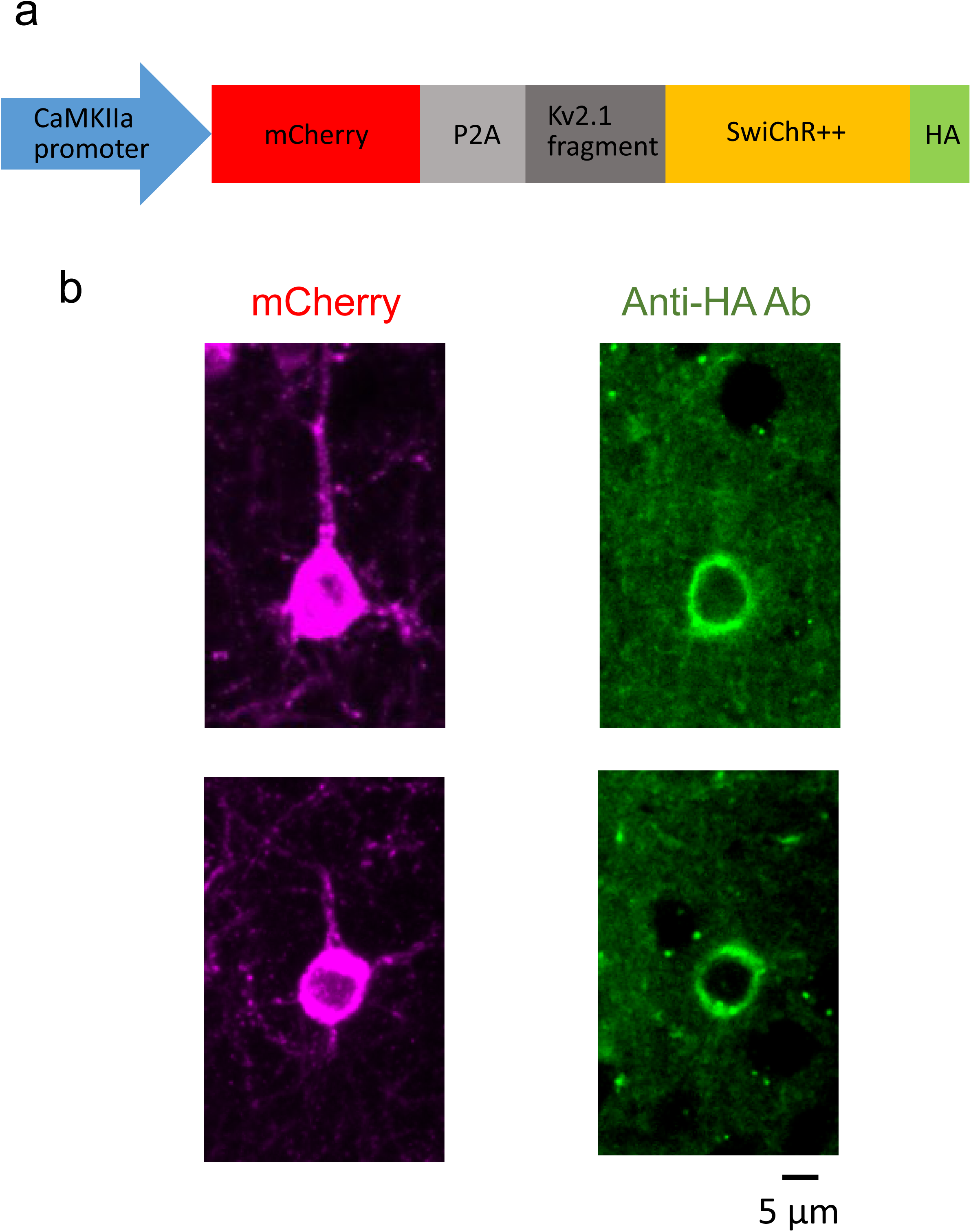
Localization of optogenetic protein at soma by using soma targeting signal. **a.** The construction of an AAV vector to express inhibitory opsin fused with influenza hemagglutinin (HA) epitope tag. **b.** Expression of mCherry (left) and immunostaining against HA using fluorescence antibody (right). The distribution of fluorescence signals indicated that inhibitory opsins are localized to cell bodies.

**Figure S3.**
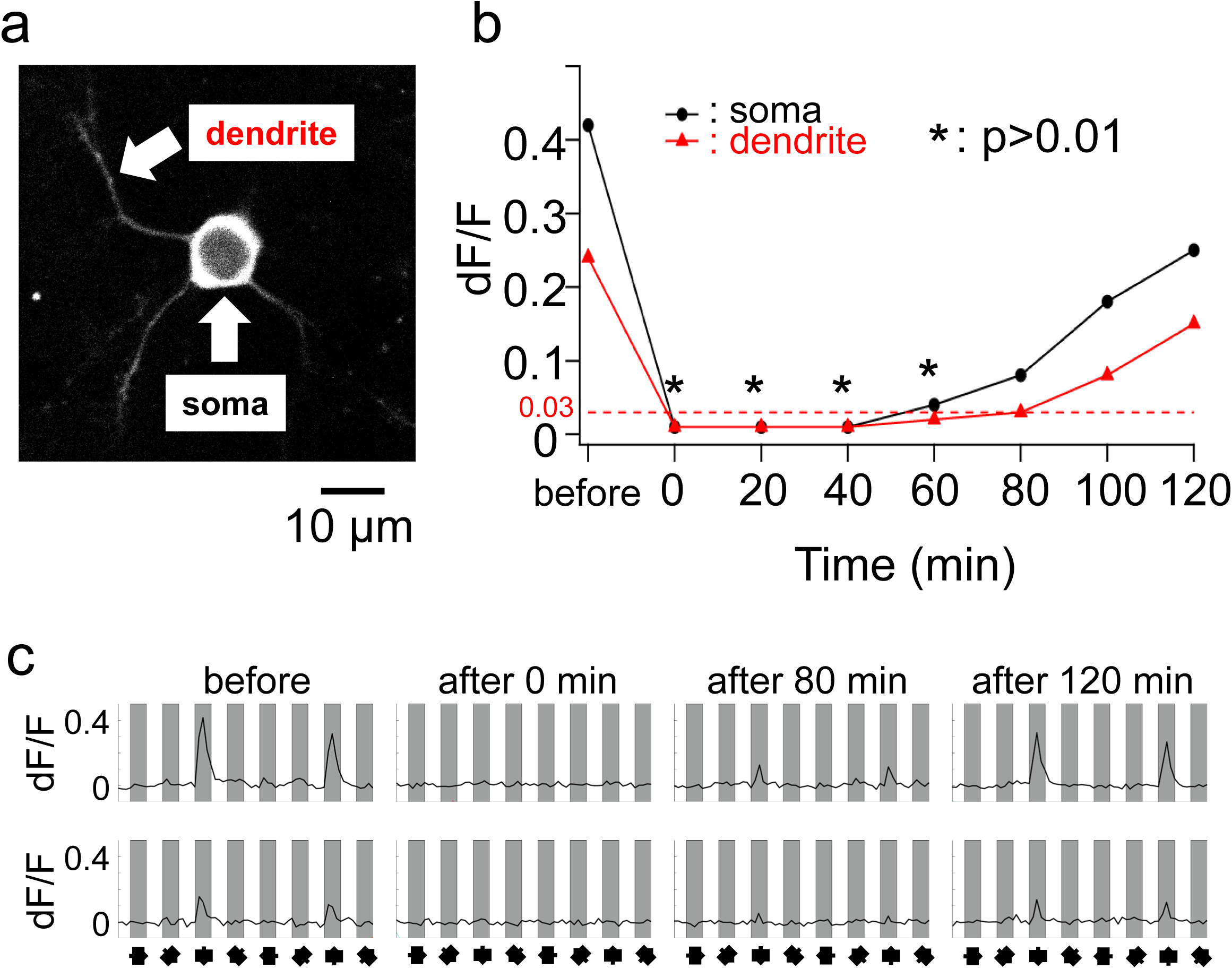
The inhibition of somatic activity by one-time photostimulation lasts for approximately an hour. **a.** GCaMP6s and SwiChR++ are coexpressed in cortical neurons, and the effect of photoinhibition of the soma is measured at the soma and dendrite during visual stimulation (the locations of measurement are shown with white arrows). **b.** Calcium signal changes at the soma and dendrite before and after photoinhibition of the soma. The asterisk means that the signal change is not significant (p=0.01 is the threshold for calculating the responsiveness, see Methods for defining the p value of the responsiveness). The loss of somatic and dendritic activity lasts for approximately an hour after photoinhibition of soma. dF/F=0.03 is the threshold of the signal change for calculating the responsiveness. **c.** The somatic (upper panels) and dendritic (lower panels) responses are shown at various time points before and after photoinhibition of the soma. Approximately 2 hours after photoinhibition, the signal change of soma and dendrite recover to ∼60% of the pre-photoinhibition values.

**Figure S4.**
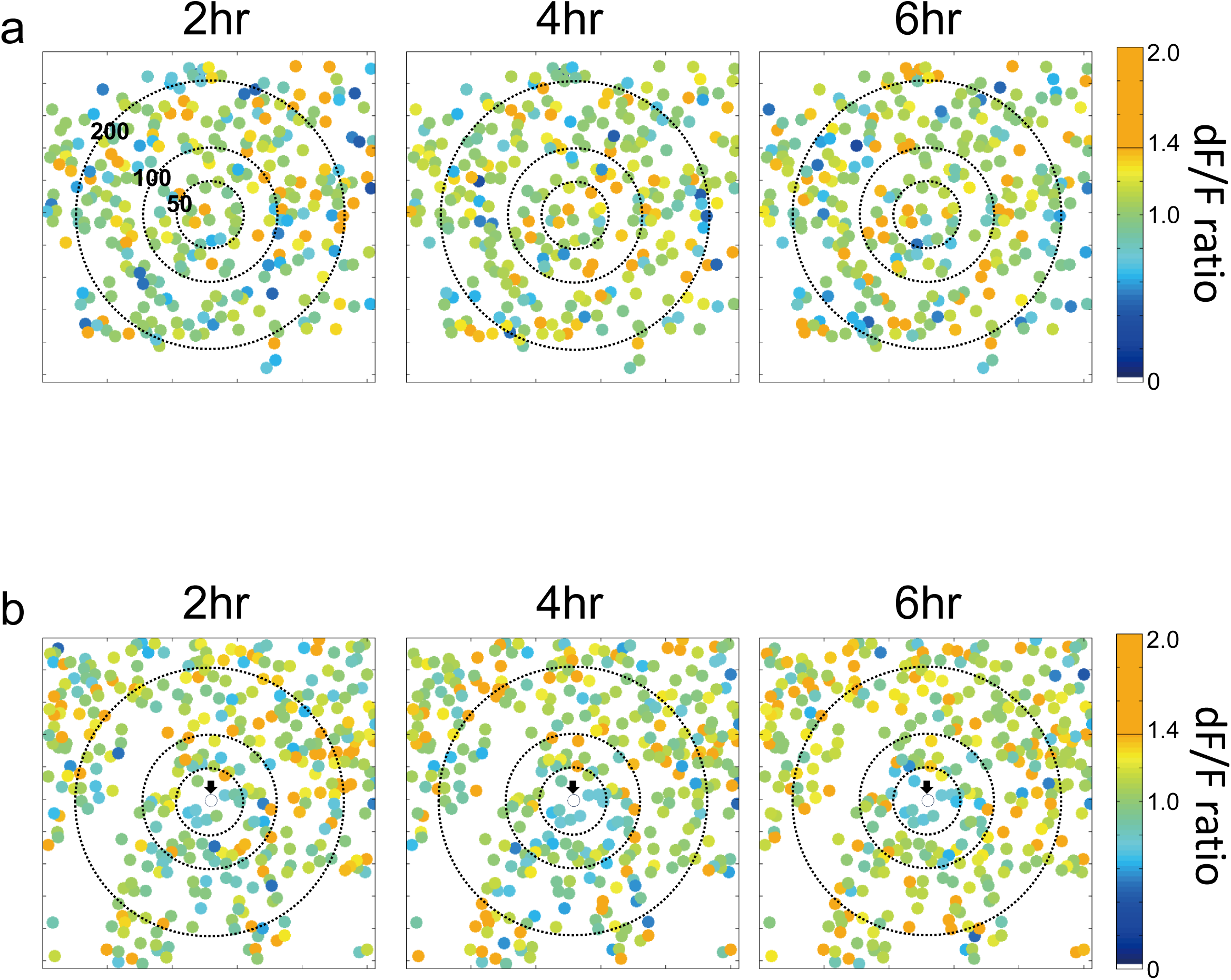
dF/F map was not largely changed after the photoinhibition. The dF/F ratio map of the imaged plane at a cortical depth of 150 µm from the pia. The dF/F ratio map is calculated by dividing the dF/F map at each time point by the dF/F map at time zero. The dF/F ratio map without photoinhibition (**a**) and with photoinhibition (**b**) are shown.

**Figure S5.**
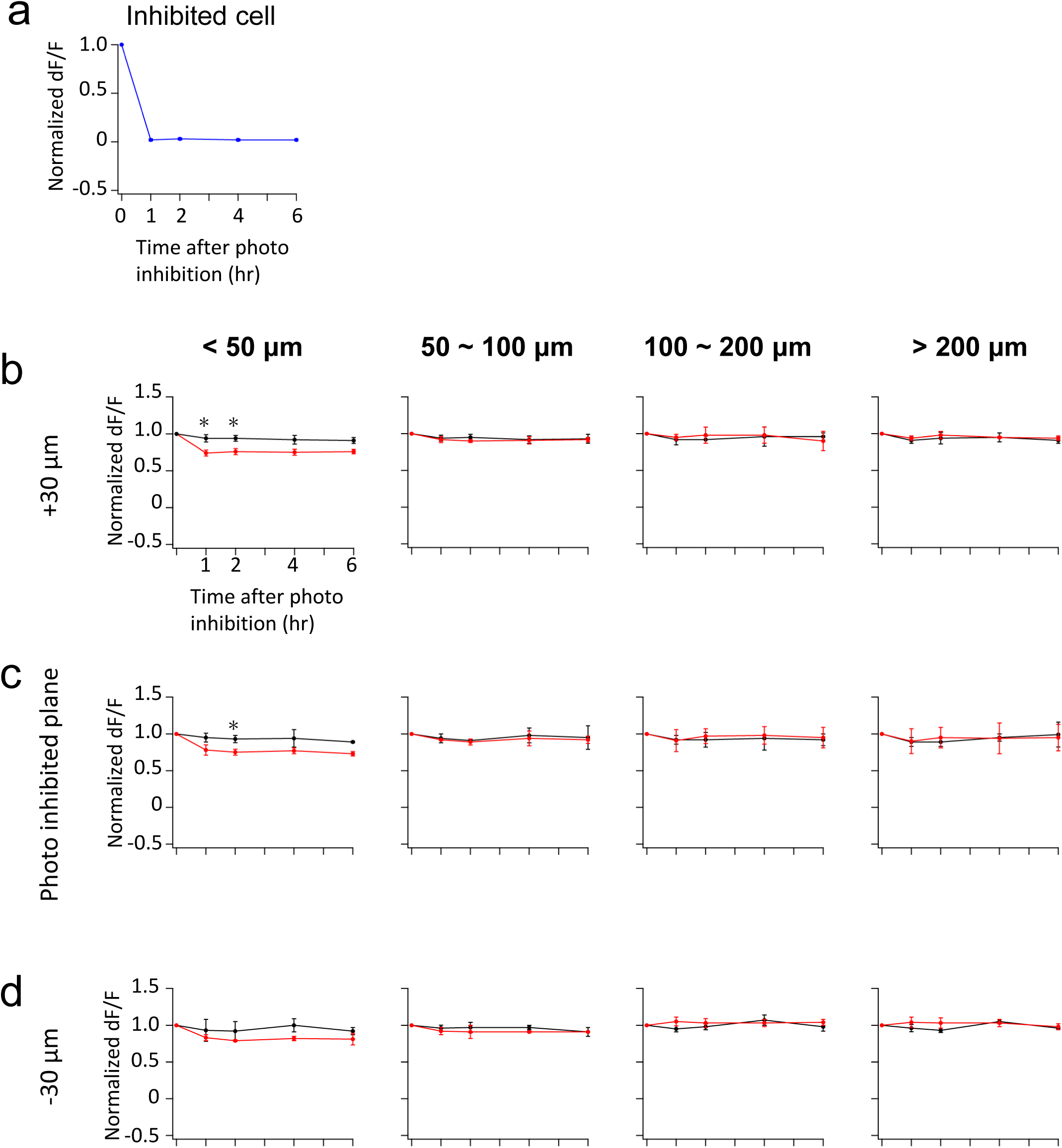
Photoinhibition did not largely change the activity of surrounding neurons. dF/F ratio map shown in Supplementary Figure 3 were quantitatively analyzed. **a.** Normalized dF/F of photoinhibited neuron (normalized to time zero) are plotted as a function of time. **b, c, d.** The normalized dF/F (normalized to time zero) of the surrounding neurons are plotted as a function of time after photoinhibition or the same time period without photoinhibition (control). Surrounding neurons are categorized by distance from the center of the FOV (no photoinhibition) or photoinhibited neuron (with photoinhibition). Normalized dF/F of surrounding neurons are calculated as mean and SD. Analyses are performed at three different depth planes; the photoinhibited plane (**c**) and planes above (**b**, +30 μm) and below (**d**, −30 μm) the photoinhibited plane. Only within 50 μm of the photoinhibited neuron, a statistically significant decrease in signal change was observed (marked with *, **b**, 1hr: p=0.022, 2hr: p=0.024; **c**, 2hr: p=0.041, unpaired-sample t-test with the Bonferroni correction).

**Figure S6.**
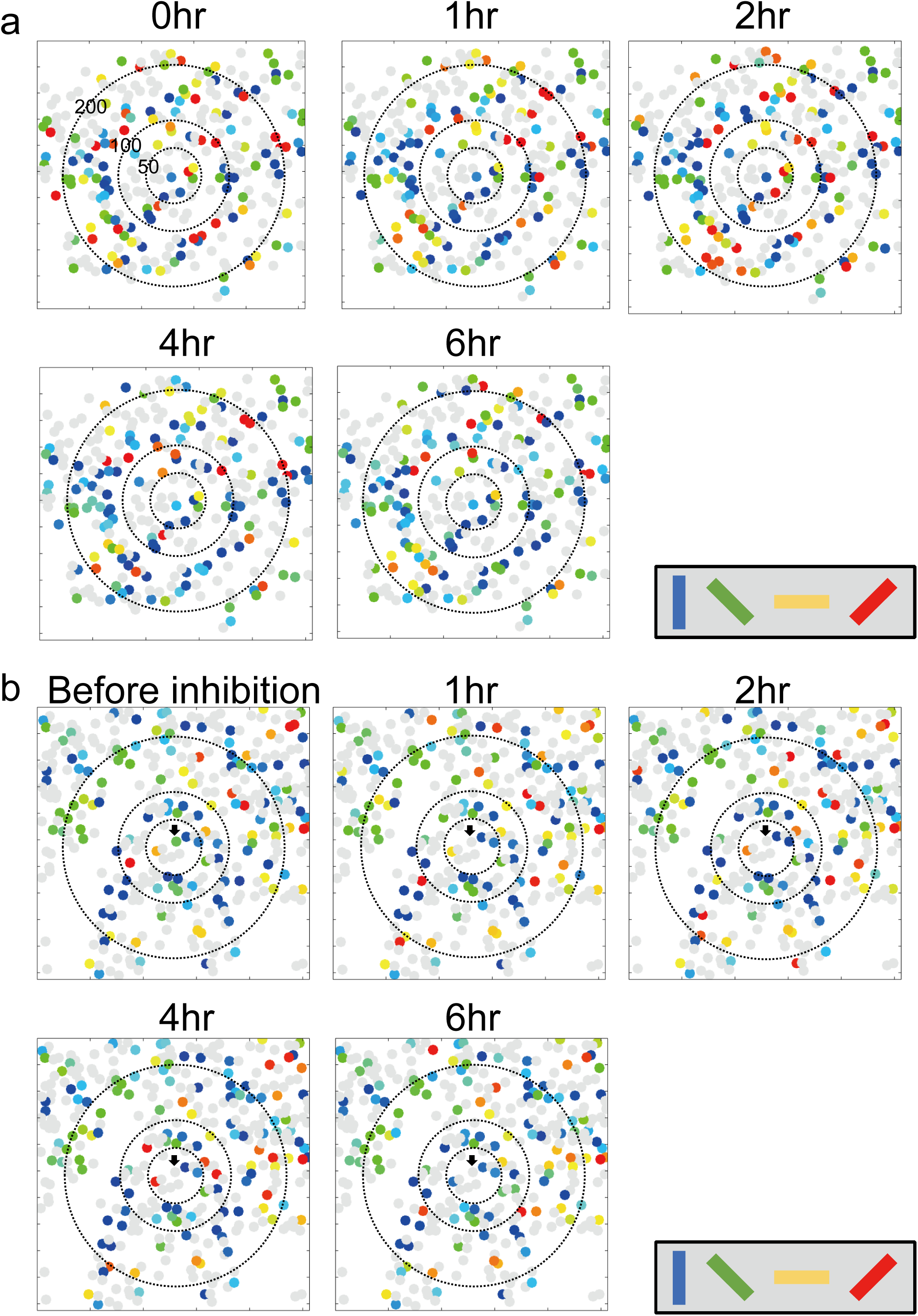
Orientation selectivity map was not changed after the photoinhibition. The orientation selectivity map of the imaged plane at a cortical depth of 150 µm from the pia. The orientation selectivity map without photoinhibition (**a**) and with photoinhibition (**b**) are shown.

**Figure S7.**
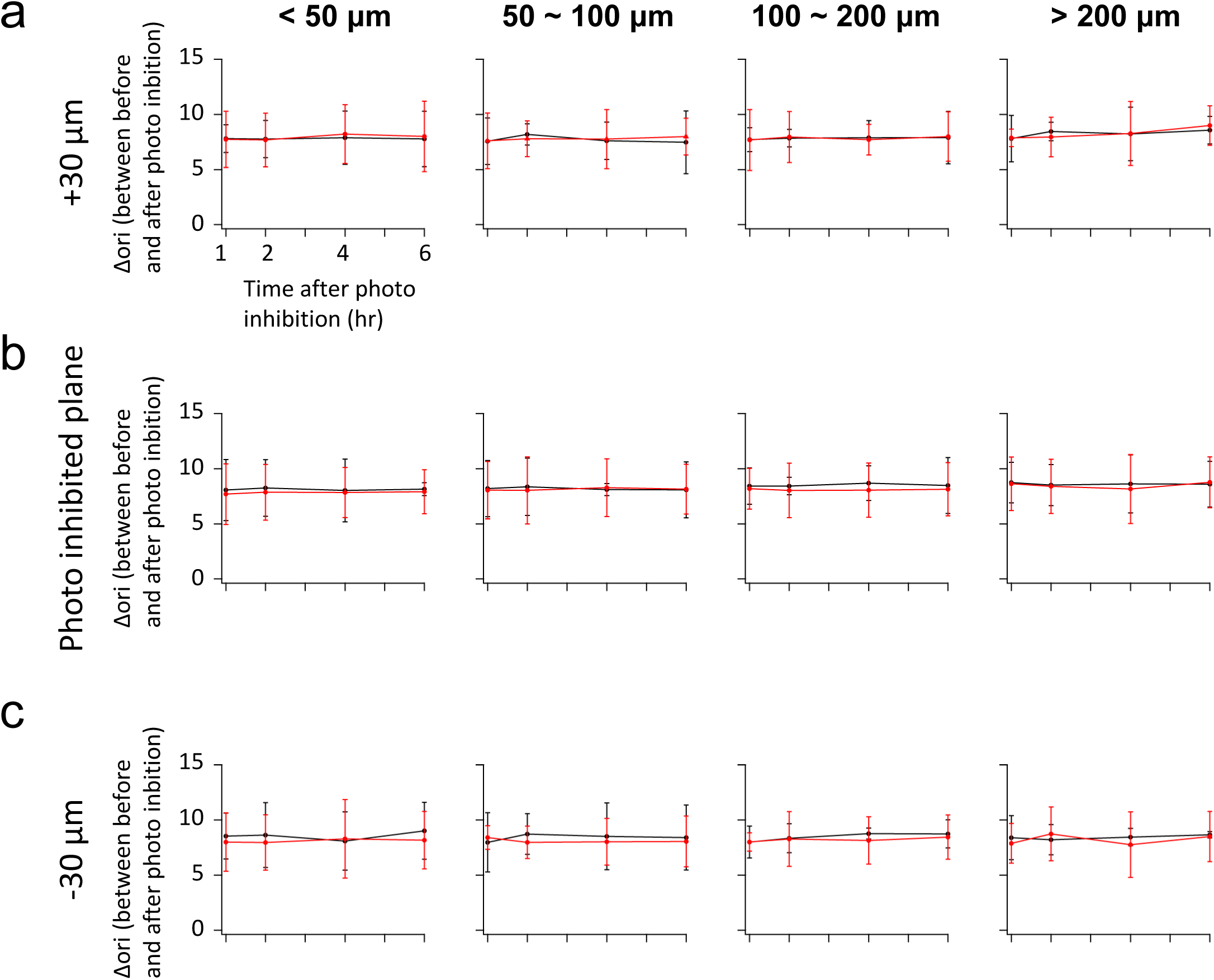
Photoinhibition did not change the orientation selectivity of the surrounding neurons. Orientation selectivity map shown in the Supplementary Figure 5 were quantitatively analyzed. The difference in the preferred orientation of the surrounding neurons is plotted as a function of time after photoinhibition or the same time period (in the case of no photoinhibition). Surrounding neurons are categorized by distance from the center of the FOV (no photoinhibition) or photoinhibited neuron (with photoinhibition). The difference in the preferred orientation of surrounding neurons is calculated as mean and SD. Analyses are performed at three different depth planes; the photoinhibited plane (**b**) and planes above (**a**, +30 μm) and below (**c**, −30 μm) the photoinhibited plane. No statistically significant difference was observed between with and without photoinhibition (unpaired-sample t-test with the Bonferroni correction).

**Figure S8.**
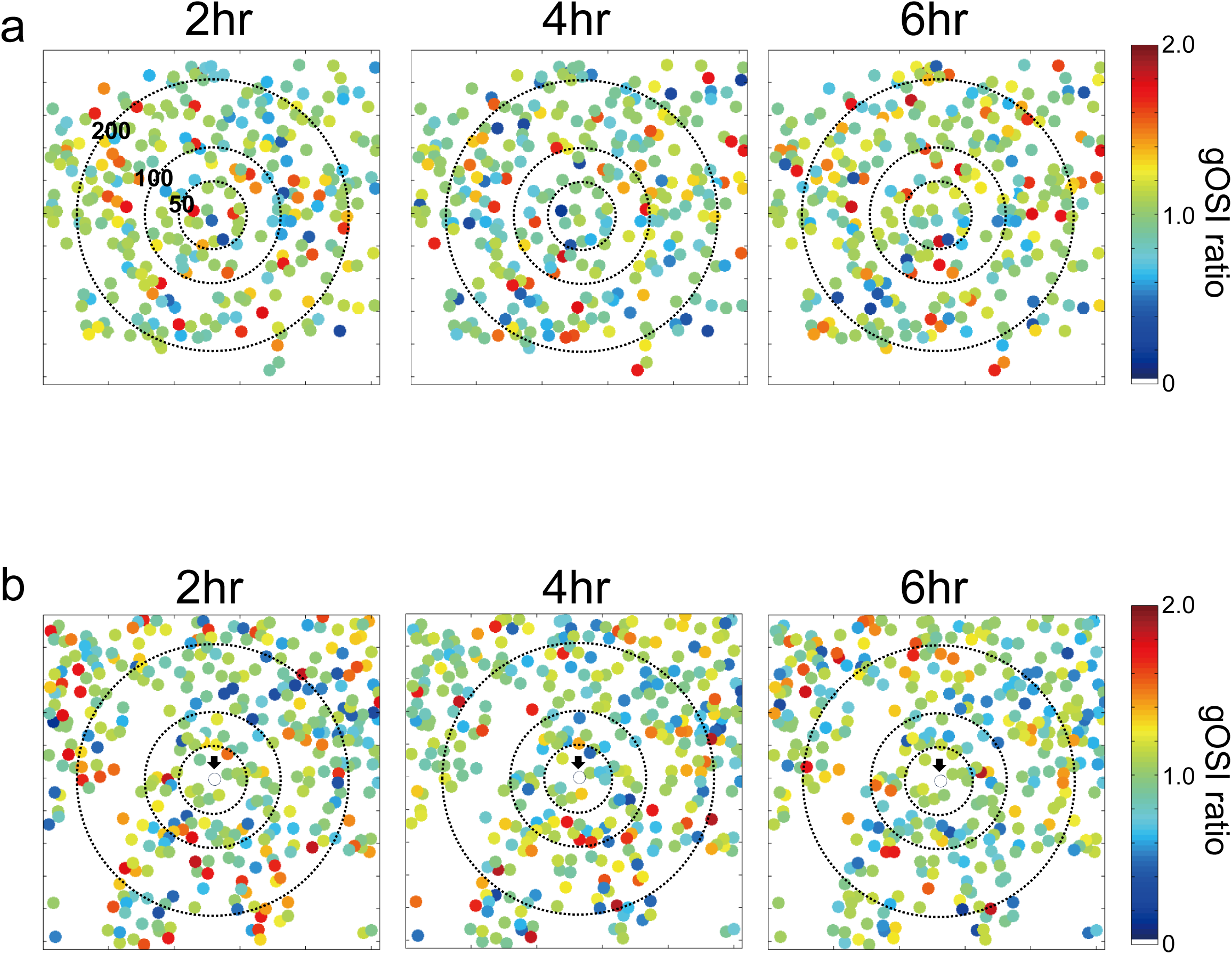
gOSI map was not changed after the photoinhibition. The gOSI ratio map of the imaged plane at a cortical depth of 150 µm from the pia. The gOSI ratio map is calculated by dividing the gOSI map at each time point by the gOSI map at time zero. The gOSI ratio map without photoinhibition (**a**) and with photoinhibition (**b**) are shown.

**Figure S9.**
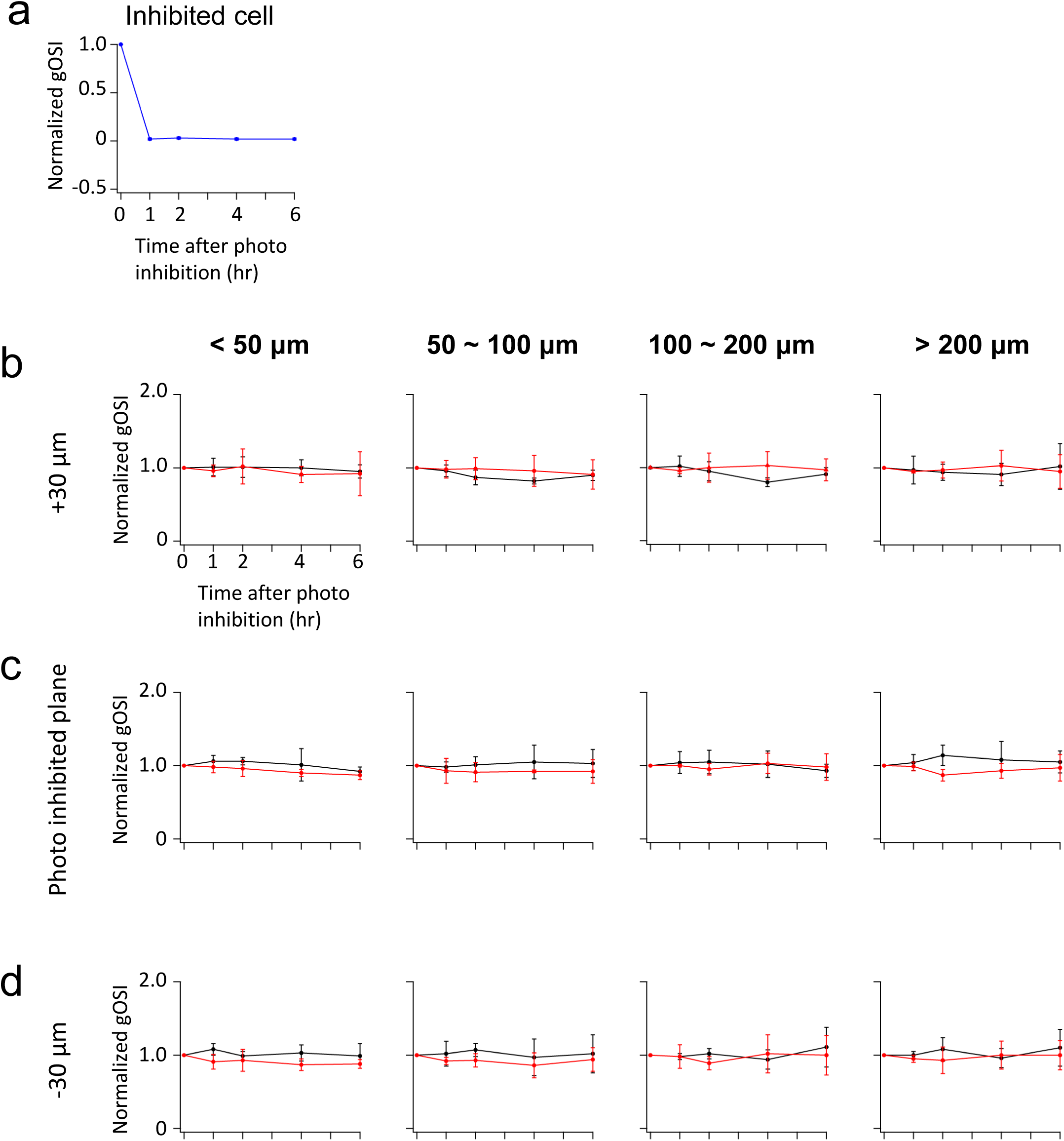
Photoinhibition did not change the gOSI of the surrounding neurons. gOSI map shown in the Supplementary Figure 7 were quantitatively analyzed. **a.** Normalized gOSI of photoinhibited neuron (normalized to time zero) are plotted as a function of time. **b, c, d.** The normalized gOSI (normalized to time zero) of the surrounding neurons is plotted as a function of time after photoinhibition or the same time period (in the case of no photoinhibition). Surrounding neurons are categorized by distance from the center of the FOV (no photoinhibition) or photoinhibited neuron (with photoinhibition). Normalized gOSI of surrounding neurons are calculated as mean and SD. Analyses are performed at three different depth planes; the photoinhibited plane (**c**) and planes above (**b**, +30 μm) and below (**d**, −30 μm) the photoinhibited plane. No statistically significant difference was observed between with and without photoinhibition (unpaired-sample t-test with the Bonferroni correction).

**Figure S10.**
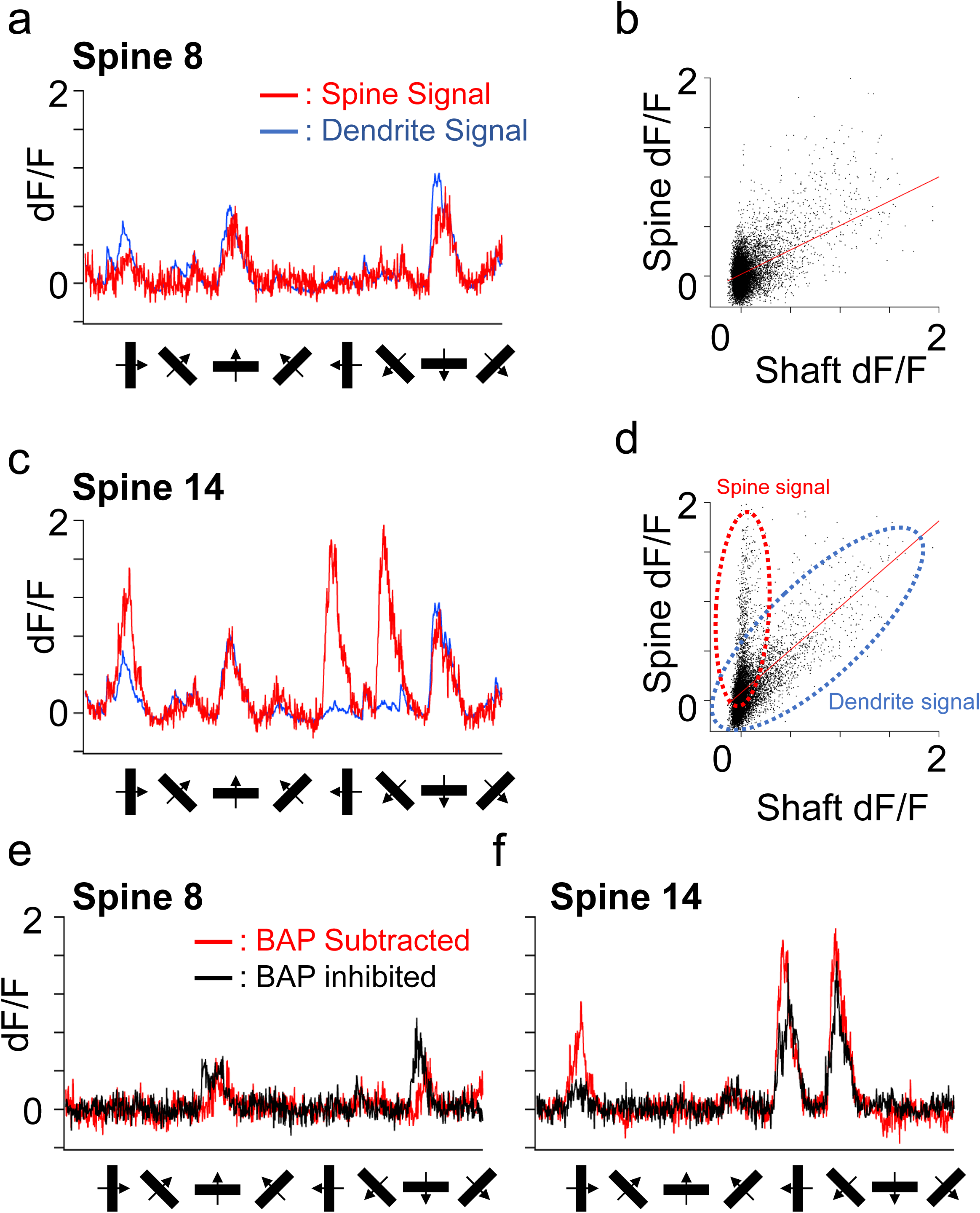
Spine-specific signal estimation by the subtraction method. **a.** An example of a spine differently tuned from the soma. The preferred orientation of the spine (red) is 0° (vertical orientation), and that of the shaft (blue) is 90° (horizontal orientation). **b.** The dendrite signal (blue dotted area) is subtracted from the spine signal (red dotted area); the correlation coefficient is calculated with robust regression (yellow line). Because of the clear separation of the spine signal component and the dendrite signal component, the estimation of the slope is accurate. **c.** An example of a spine similarly tuned as soma. The preferred orientations of the spine and dendrite are matched at 90° (horizontal orientation). **d.** The dendrite signal (blue dotted area) is subtracted from the spine signal (red dotted area); the correlation coefficient is calculated by robust regression (yellow line). Because of the overlap of the spine signal component with the dendrite signal component, the estimation of the slope is inaccurate. **e**. **f**. The resulting spine time course of the BAP subtracted method and the same spine time course of our new photoinhibited method are compared. Two methods showed the similar time course for both spines whose preferred orientation was the same as the soma (**e**) and different from the soma (**f**).

**Figure S11.**
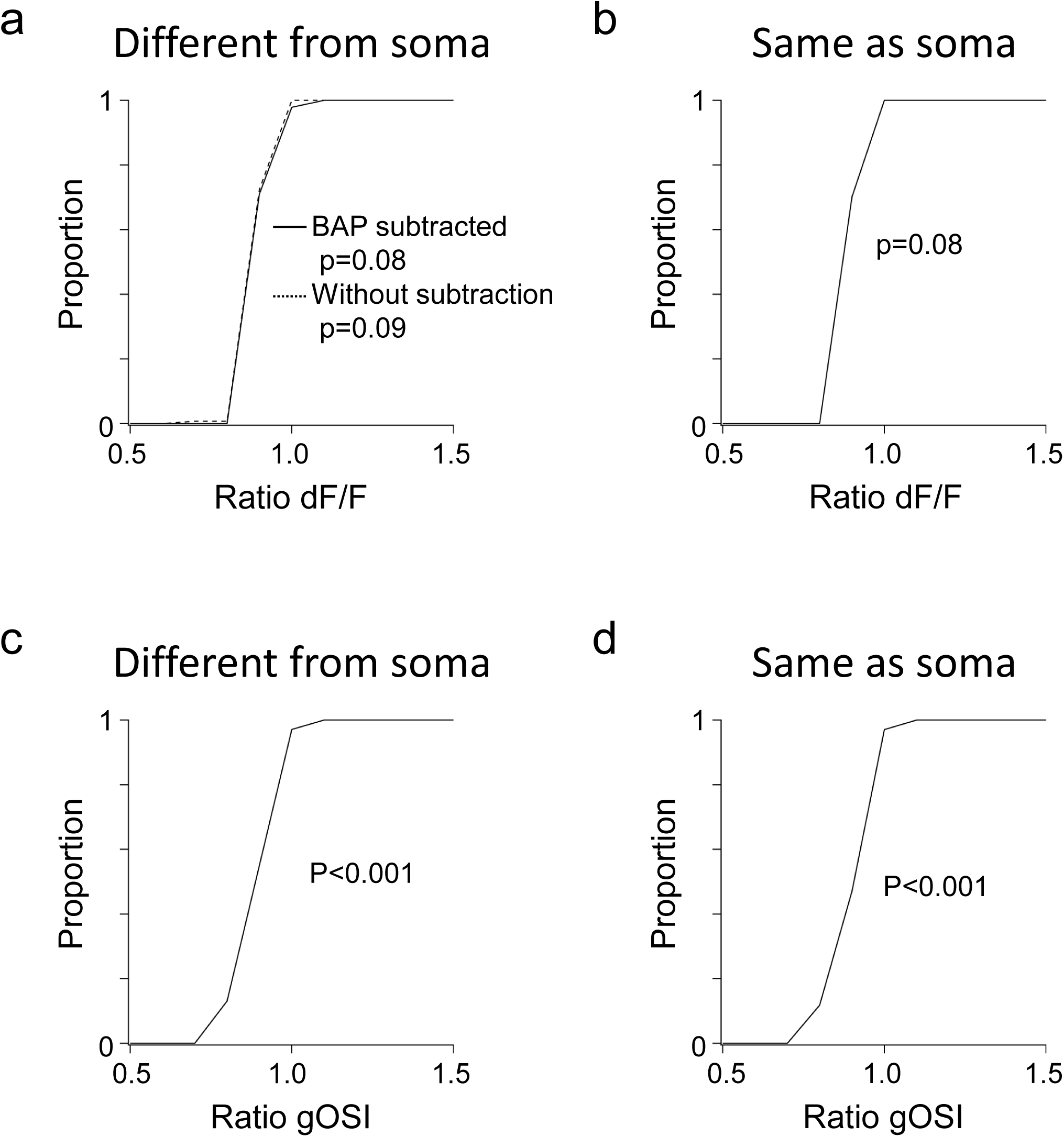
Comparison between the subtraction and photoinhibition methods for the visually responsive spine activity. dF/F values obtained by without-BAP-subtraction, BAP-subtraction or BAP-inhibition methods or gOSI values by BAP-subtraction and BAP-inhibition methods shown in Figure 2 were divided and calculated the ratios. **a**. Cumulative plots of the dF/F ratio of all the orientation-selective spines with preferred orientation different from soma. There was no statistically significant difference (p=0.08, BAP-subtraction vs. BAP-inhibition methods; p=0.09, without subtraction vs. BAP-inhibition method; one-sample t-test). **b**. Cumulative plots of the dF/F ratio of all the orientation-selective spines with preferred orientation same as soma. There is no statistically significant difference (p=0.08, BAP-subtraction vs. BAP-inhibition methods; one-sample t-test). **c, d**. Cumulative plots of the gOSI ratio of all the orientation-selective spines with preferred orientation different from soma (**c**) and same as soma (**d**). There is no statistically significant difference in both cases (**c**, p<0.001 one-sample t-test; **d**, p<0.001 one-sample t-test).

**Figure S12.**
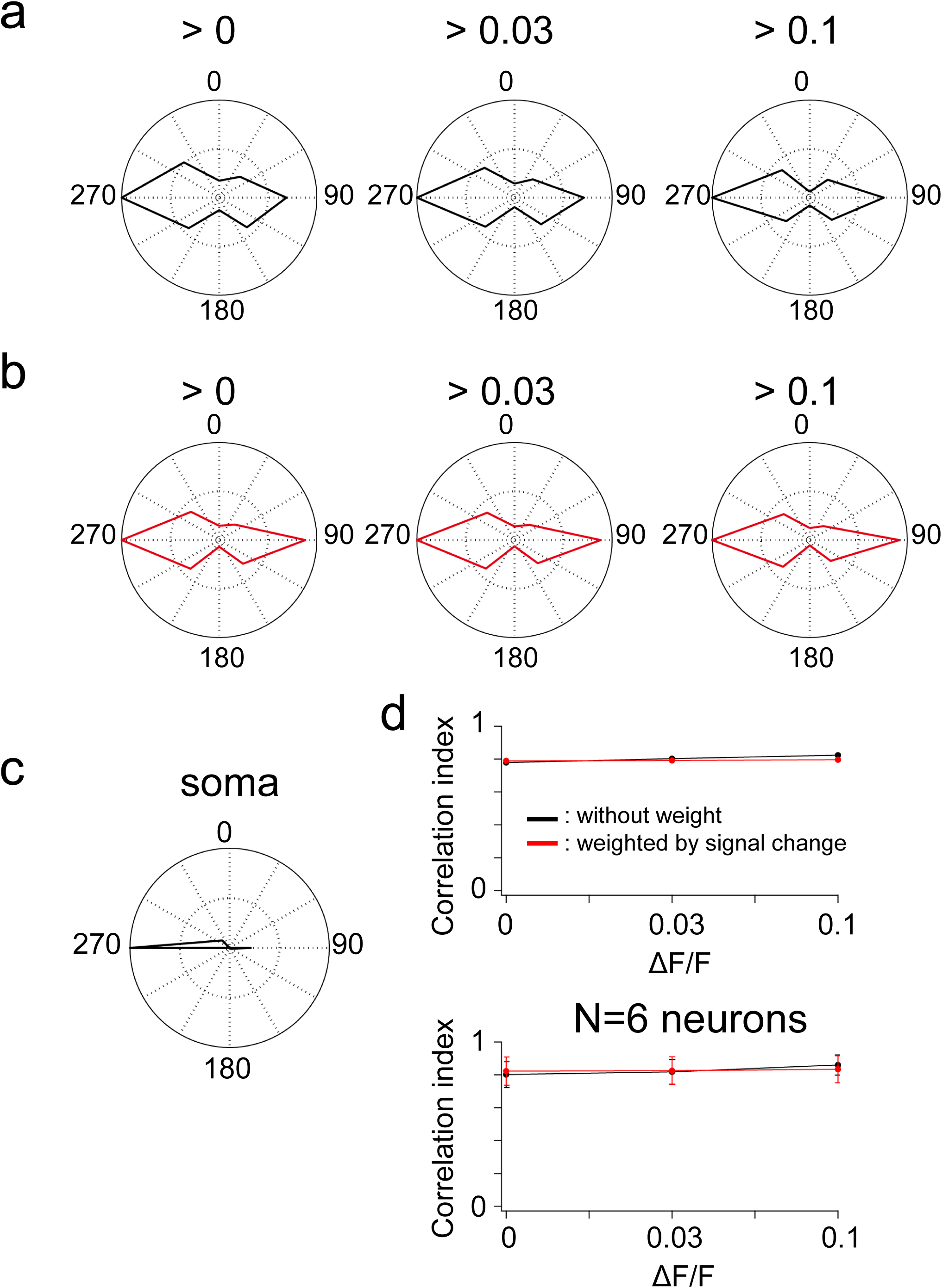
The prediction of soma tuning is not improved by considering only spines with large calcium signal changes. **a, b.** Thresholds for spine signal change are set (>0, >0.03 and >0.1) and either the number of responsive spines (**a**) or the number of responsive spines weighted by signal change (**b**) is summed at each threshold. **c.** Actual soma tuning. **d.** CIs are plotted as a function of the threshold for a single neuron (upper panel) and the population average (lower panel).

**Figure S13.**
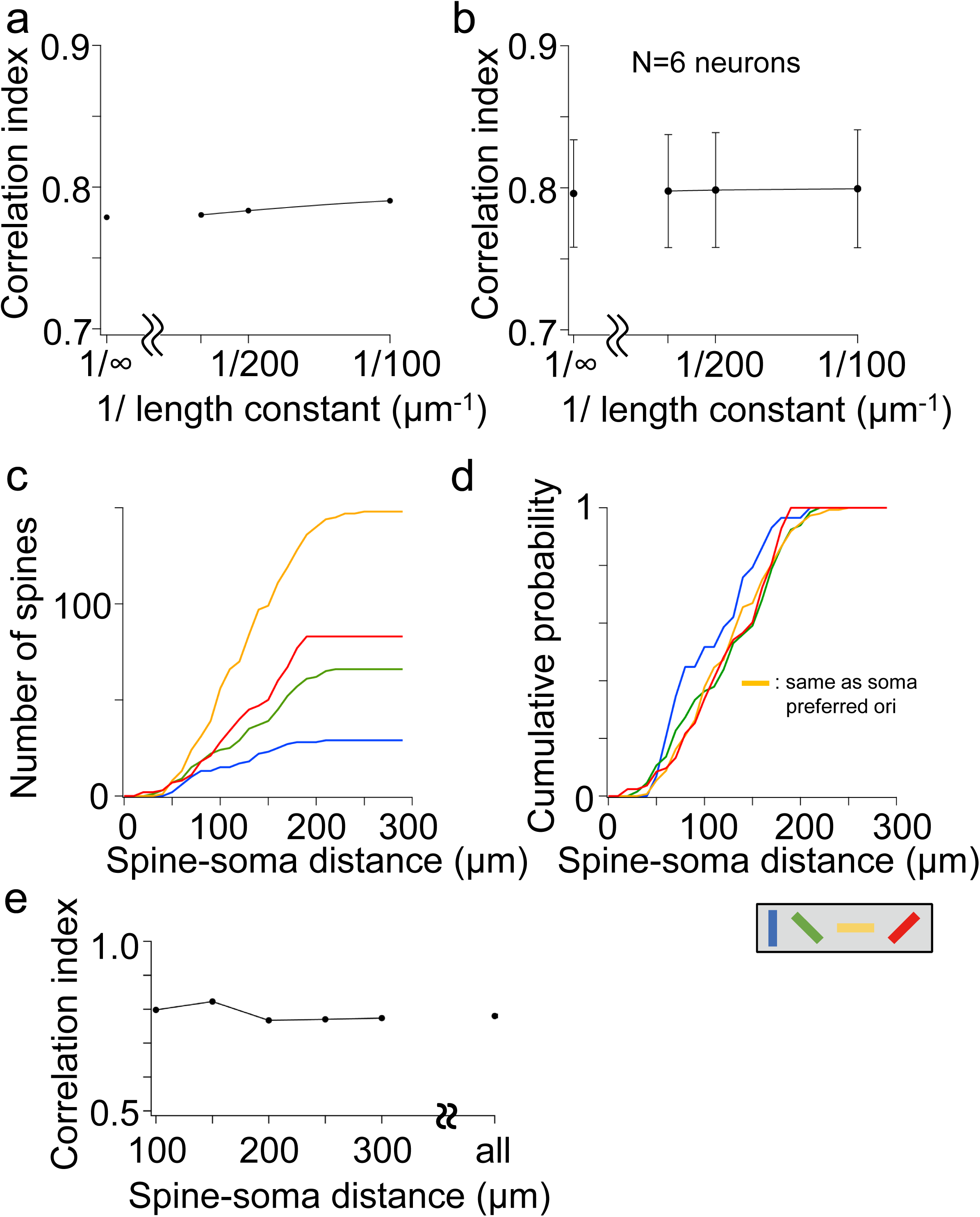
The prediction of soma tuning is not improved by considering the distance between the spines and the soma. **a.** An example of single neuron. **b.** Population averages from 6 neurons. **c.** The number of spines with various preferred orientations is plotted as a function of the spine-soma distance. **d.** The distribution profile of spines of each preferred orientation. **e.** CIs between the actual soma tuning and the predicted soma tuning using spines located in the limited distance from soma (100 ∼ 300 μm from the soma) or all spines are plotted.

**Figure S14.**
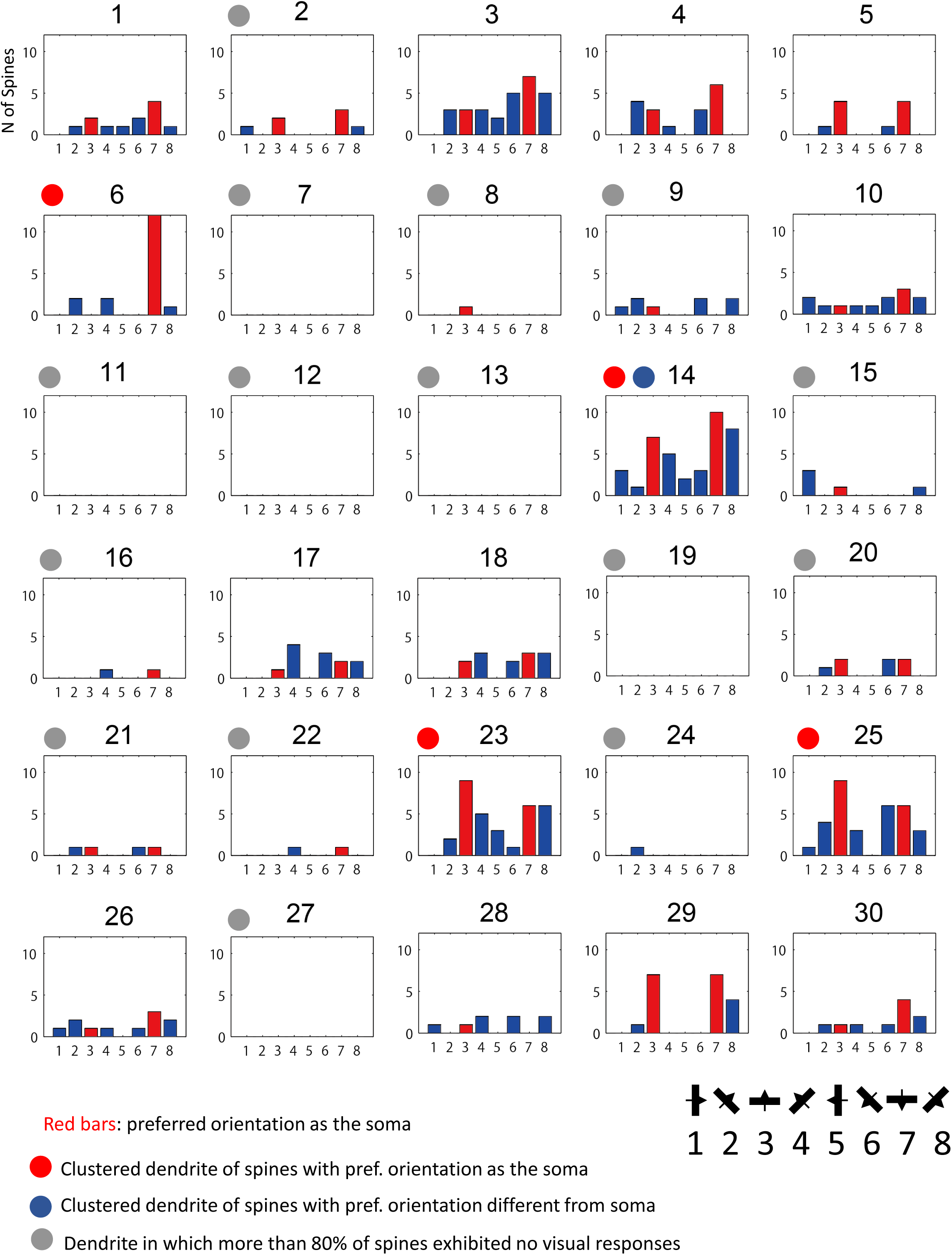
Functional clustering of spines on the dendritic segment. The histograms of number of spines that responded to any orientations are shown for each dendritic segment (neuron 6 in Table 1). Red bars represent the same orientation as the preferred orientation of the soma.

**Figure S15.**
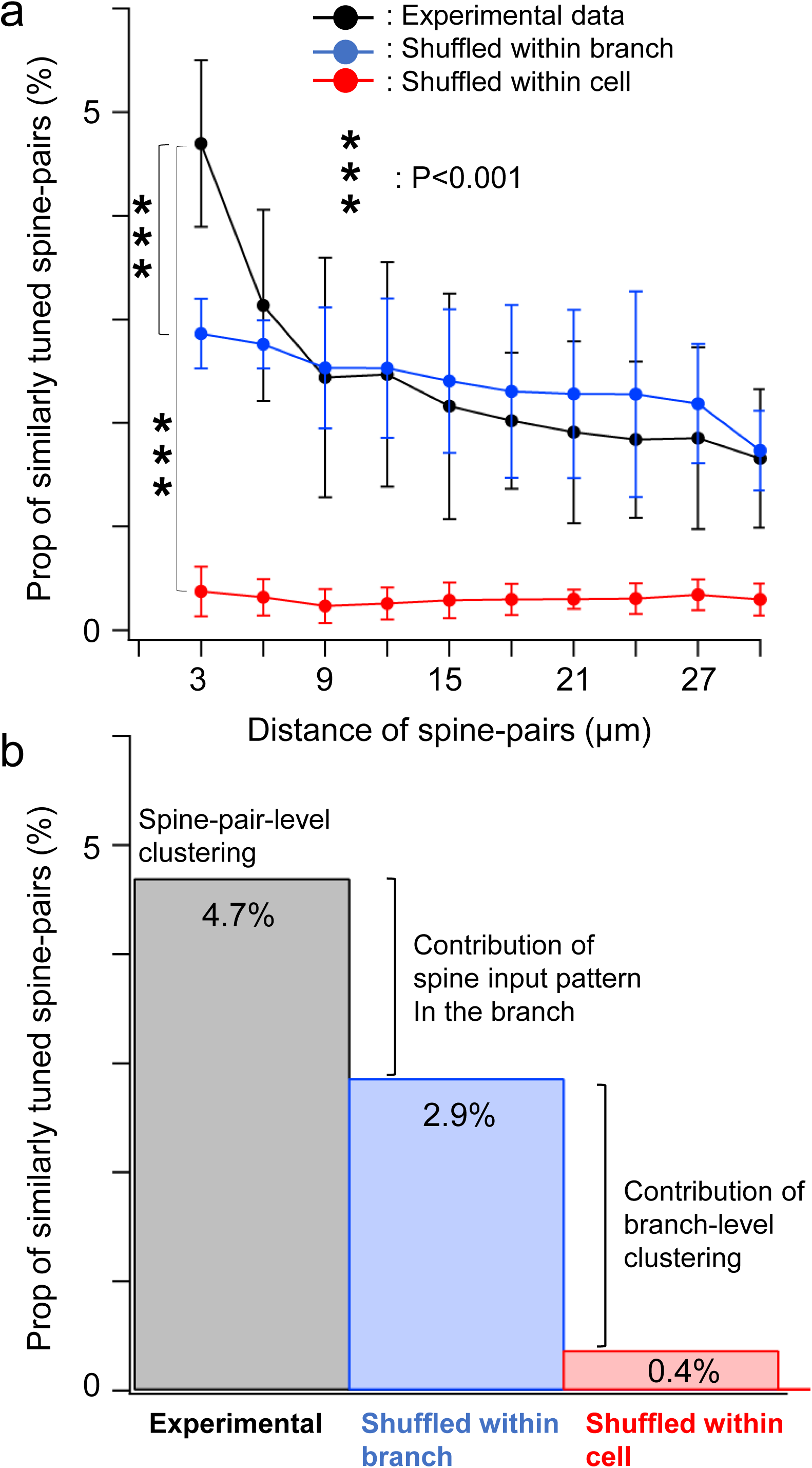
Relationship between two forms of functional clustering. To investigate the relationship between segment-level clustering and spine-pair-level clustering, we evaluated the contribution of segment-level clustering to spine-pair-level clustering. **a**. We calculated the proportion of similarly tuned spine-pairs to all spine-pairs at different distances of paired spines (30 μm length in 3 μm steps) on each segment and compared it between the experimental data (black lines) and the randomized data (blue and red lines). Randomization was performed among all spines either within a segment (blue line) or across cell (red line). Statistically significant difference was observed only at the spatial distance of 3 μm (p=0.004, experimental vs. shuffled within segment; p<0.001, experimental vs. shuffled across cell; Mann–Whitney U-test with the Bonferroni correction). **b**. The contribution of segment-level clustering to the spine-pair-level clustering was evaluated. Randomizing within a segment significantly reduced the proportion of similarly tuned spine pairs (the difference in height between black and blue bars), and this reduction (1.8%) correspond to the contribution of spine input pattern within the clustered segment. Randomizing across cell further significantly reduced the proportion of similarly tuned spine pairs (the difference in height between blue and red bars), and this reduction (2.5%) may correspond to the contribution of the segment level clustering. Thus, it can be estimated that segment-level clustering may contribute about half to the spine-pair-level clustering.

**Figure S16.**
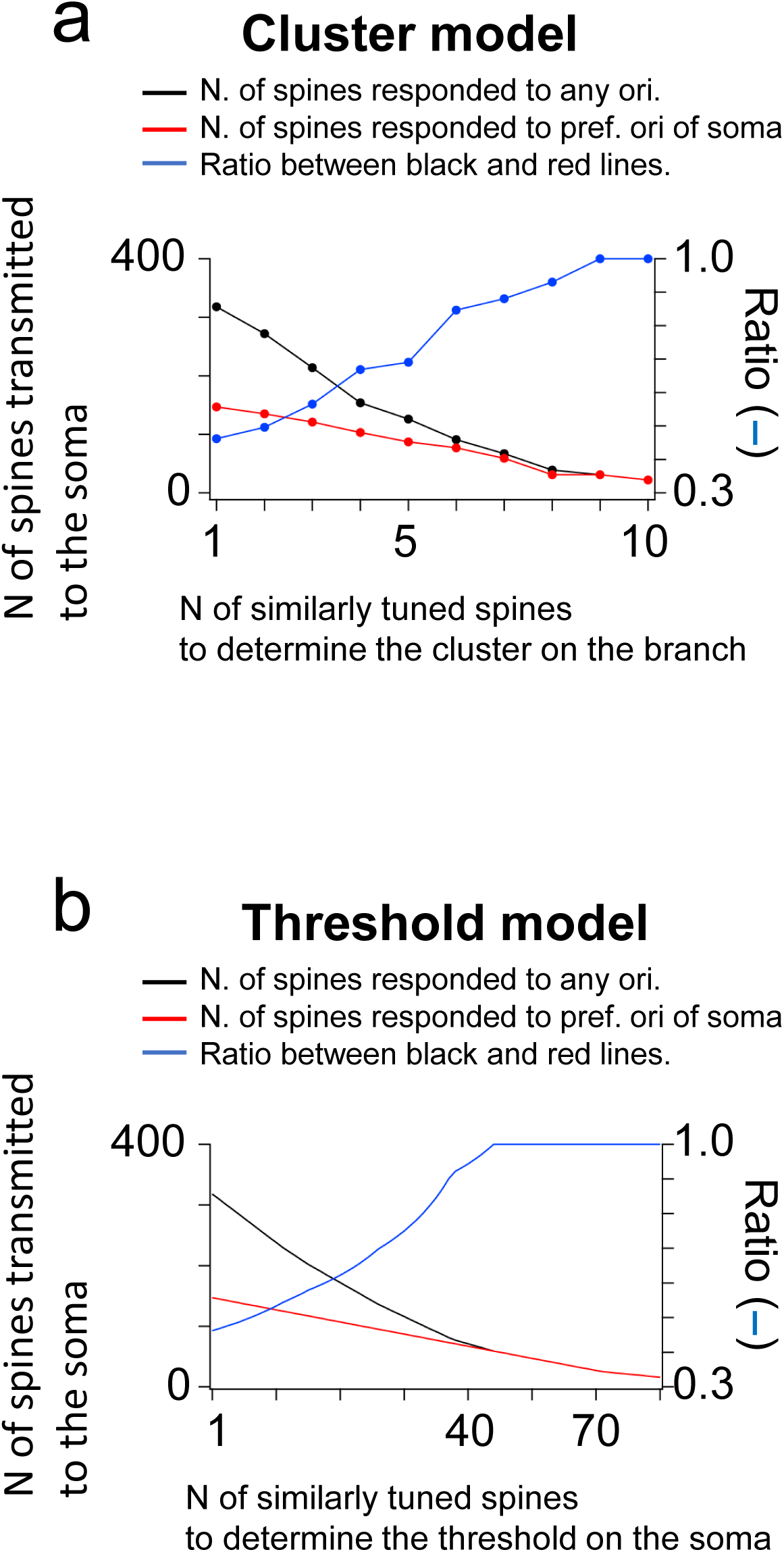
Summation of the number of responsive spines, with or without weighting, cannot reproduce the somatic tuning curve. **a.** The number of responsive spines is plotted as a function of the number of spines in a cluster. The black line shows number of spines responding to any orientation, and the red line shows number of spines responsive to the preferred orientation of the soma. The blue line represents the ratio between the red and black lines. **b.** The number of responsive spines is plotted as a function of the number of spines for the threshold. Black line shows number of spines responding to any orientation, and red line shows number of spines responsive to the preferred orientation of the soma. The blue line represents the ratio between the red and black lines.

**Figure S17.**
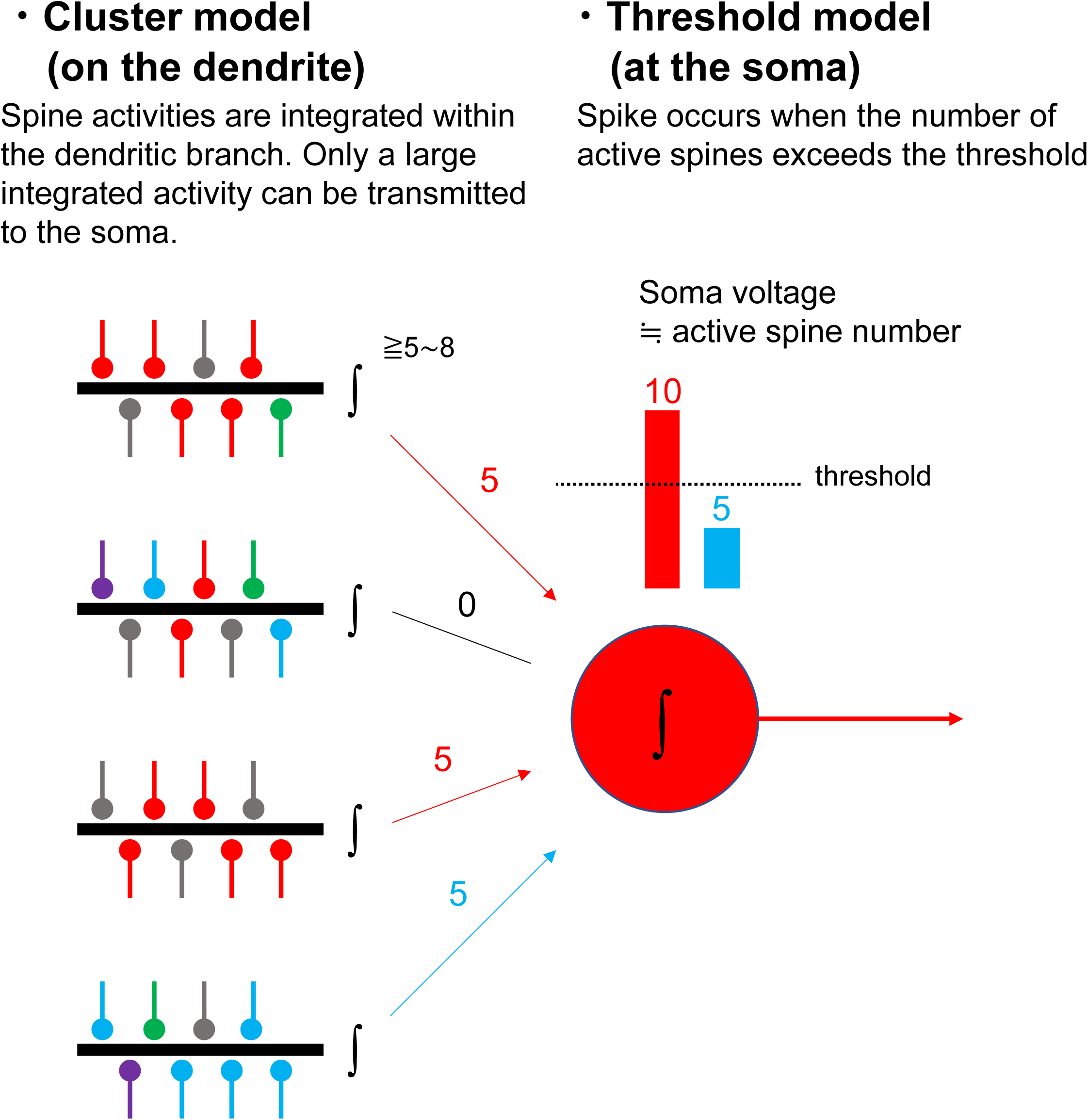
Combination of cluster and threshold models for integration of synaptic inputs. Our results indicate that synaptic integration can be explained by a combination of cluster and threshold models. The cluster model is a nonlinear integration of synaptic inputs that occur at dendrites with clusters of simultaneously responding spines. Effective integration can occur from clusters with more than 4−6 spines, and this integrated response can be transmitted to the soma. The threshold model works at the soma to determine when the soma spikes. Assuming that the somatic membrane voltage is proportional to the number of active spines whose signals are transmitted to the soma, the soma spikes when the number of active spines exceeds the threshold. Although either the cluster model or the threshold model alone can approximately predict the soma tuning, the combined model can better reproduce the soma tuning curve.

